# Accurate and efficient phylogenetic inference through end-to-end deep learning

**DOI:** 10.1101/2025.09.30.679045

**Authors:** Xinru Zhang, Shizhe Ding, Chungong Yu, Jianquan Zhao, Dongbo Bu

**Author notes:** Correspondence: Correspondence should be addressed to Shizhe Ding or Dongbo Bu.

## Abstract

Accurate phylogenetic inference is crucial for understanding evolutionary relationships among species. Deep learning technique has been introduced for phylogenetic inference; however, the existing deep learning-based approaches either suffer from limited accuracy as they split inference into several disjoint stages, or exhibit low efficiency and hardly apply to the cases with over 20 species. Here we present an accurate and efficient approach to phylogenetic inference. Our approach, called NeuralNJ, employs an end-to-end framework that directly constructs phylogenetic trees from the input taxa, thus effectively avoiding the inaccuracy incurred by the split inference stages. The key innovation of NeuralNJ lies in its learnable neighbor joining mechanism, which iteratively joins neighbors guided by learned priority scores and thereby achieves accurate tree reconstruction. The inference accuracy is further enhanced through incorporating reinforcement learning-based tree search. Using both simulated and empirical data as representatives, we demonstrate that NeuralNJ can effectively infer phylogenetic tree with improved computational efficiency and reconstruction accuracy. The study paves the way to accurate and efficient phylogenetic inference for hundreds of taxa in complex evolutionary scenarios.

## Introduction

Phylogenetic inference aims to reconstruct evolutionary relationship among organisms through analysing their genetic information, especially molecular sequences (Brocchieri, 2001). This analytical approach has become fundamental to a variety of biological tasks, including systematics (Rannala and Yang, 2008), molecular evolution (Shao et al., 2021; Lythgoe et al., 2021; Hoyt et al., 2022) and molecular ecology (Pickering et al., 2022; Gómez-Corrales and Prada, 2020). For example, phylogenetic analysis of COVID-19 strains provided invaluable insight into their evolutionary history, thus greatly facilitating timely evidence-based public health decisions (Morel et al., 2020; Li et al., 2021). Despite its great importance, phylogenetic inference is challenged as the number of possible tree topologies grows super-exponentially with the number of species (Felsenstein, 1978; Chor and Tuller, 2005). Thus, developing accurate and efficient phylogenetic inference approaches is highly desired.

Classical phylogenetic inference approaches, which rely on expert-designed heuristics to construct phylogenetic trees, can be divided into two categories, namely, distance-based and character-based approaches. The distance-based approaches first estimate pairwise distances among the given species, and then construct a phylogenetic tree that optimally satisfies the estimated distance constraints through clustering or neighborjoining (Jukes et al., 1969; Fitch and Margoliash, 1967; Sokal and Michener, 1958; Saitou and Nei, 1987). Unlike the distance-based approaches using a single number to summarize the difference between two species, the character-based approaches explore full information of organisms’ characters (traits, features, or DNA sequences) to find the most plausible phylogenetic tree explaining the character distribution. The widely-used explaining principles include maximum parsimony, which builds a tree with the minimized character changes (Farris, 1970; Swofford and Sullivan, 2009), maximum likelihood (ML) under a specific evolutionary model (Felsenstein, 1981; Kozlov et al., 2019; Nguyen et al., 2014; Minh et al., 2020; Guindon et al., 2010), and Bayesian inference that identifies the tree with the maximum posterior probability (Felsenstein, 2004; Price et al., 2010; Bouckaert et al., 2014; Ronquist et al., 2012). The phylogenetic tree can be constructed either through tree building (such as neighbor joining) or through iterative improving a tree using rearrangement operations like subtree pruning and regrafting (SPR) (Bordewich and Semple, 2005) and nearest neighbor interchange (NNI) (Waterman and Smith, 1978; Desper and Gascuel, 2002).

Recently, machine learning, especially deep learning, has emerged as a promising paradigm for phylogenetic inference. These approaches learn key elements of phylogenetic inference directly from training data, specifically elements such as pairwise distance (Nesterenko et al., 2025), tree rearrangement strategies (Azouri et al., 2021, 2024), and tree likelihood estimation (Xie and Zhang, 2023; Zhang, 2023). Generative adversarial network has also been incorporated into phylogenetic inference, which attempts to yield a phylogenetic tree that generates multisequence alignment (MSA) indistinguishable from the input MSA (Smith and Hahn, 2023).

The machine learning-based approaches, although exhibiting promising performance, still suffer from several limitations: *i*) Some approaches, such as PhyDL (Zhou et al., 2023) and PhyloGame (Azouri et al., 2024), focus on quartet molecular phylogeny (Ranwez and Gascuel, 2001) or small datasets with at most 20 taxa, thus restricting themselves from larger phylogenetic inference tasks. *ii*) Other approaches, such as VBPI-GNN (Zhang, 2023), ARTree (Xie and Zhang, 2023), focus on the accuracy of posterior likelihood of phylogenetic trees but pay little attention to the efficiency of tree construction. *iii*) Only partial modules of the inference procedure are trained, e.g., Phyloformer (Nesterenko et al., 2025) trains the distance estimating module only and simply uses the traditional distance-based tree construction module, thus making the training errors cannot be sufficiently explored to optimize all modules.

Here we present an accurate and efficient approach (called NeuralNJ) to phylogenetic inference which constructs the tree directly from genome sequences. Our approach adopts an end-to-end framework that propagates the training loss from the final layer back to the input layer, thus enabling optimizing all intermediate modules along the propagation and improving accuracy of phylogenetic inference. In addition, rather than learning pairwise distances between species, which is dependent on accurate distance estimation and sensitive to branch length variations, our approach learns to rank candidate subtrees using a learning-to-rank strategy, potentially offering advantages in both phylogenetic inference stability and accuracy. Our approach constructs phylogenetic trees in just one pass rather than multiple iterative steps for refinements, thus gaining the advantages of high efficiency. We evaluated NeuralNJ on simulated and empirical datasets to highlight its special features in both accuracy and efficiency. The source code of NeuralNJ, together with the datasets for training and evaluation, are publicly available at https://github.com/ZhangXinru99/NeuralNJ.

## Results

### Overview of NeuralNJ algorithm

We describe the main steps and operations of NeuralNJ in Figure 1. NeuralNJ takes genome sequences in the form of MSA as its input and constructs a phylogenetic tree using an encoder-decoder architecture that consists of a sequence encoder and a tree decoder. The details of these two modules are described as follows:

**Figure 1.**
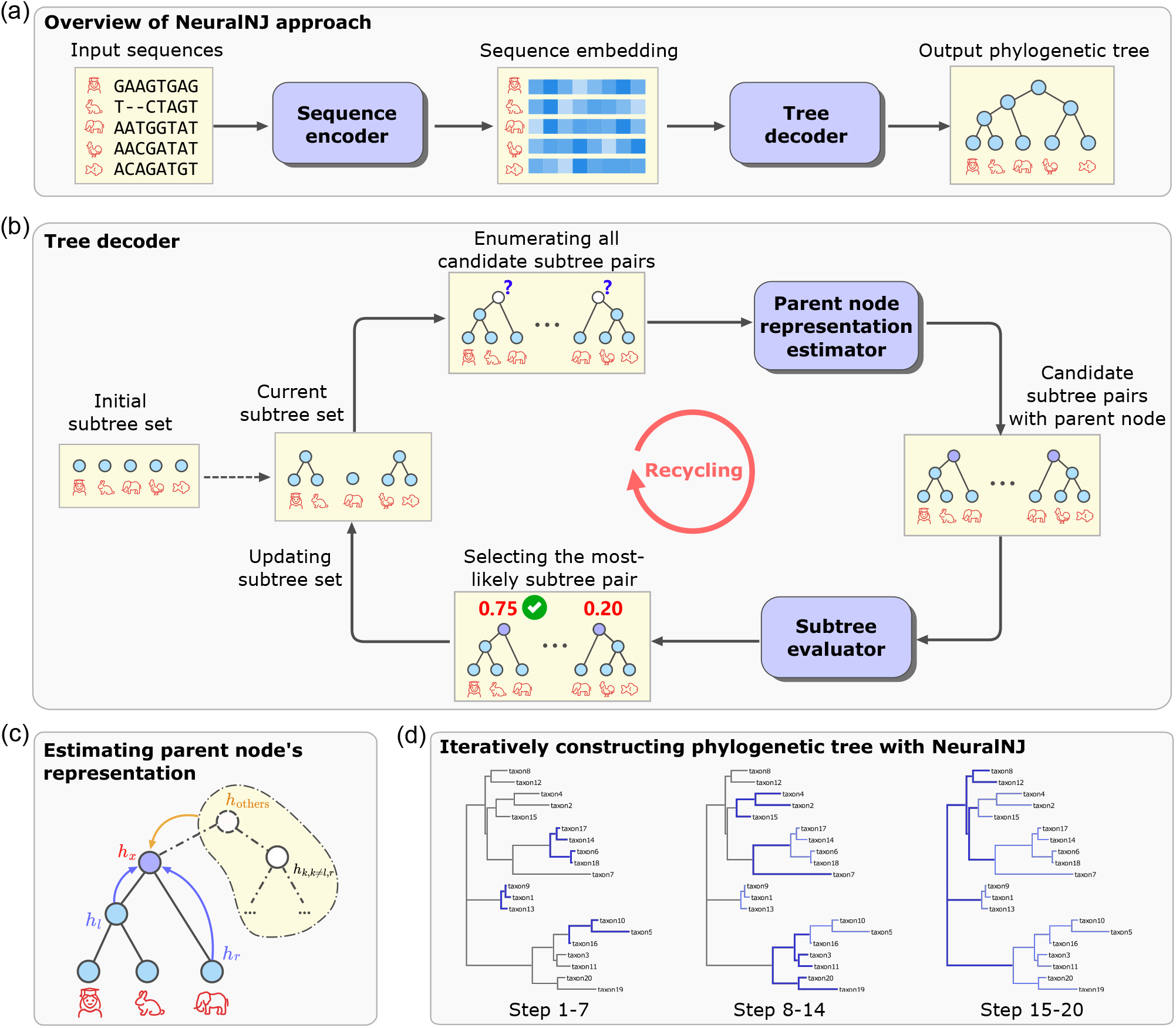
Overview of NeuralNJ architecture and workflow. (a) The overall architecture of NeuralNJ, consisting of a sequence encoder that transforms input sequences into embeddings and a tree decoder that constructs the phylogenetic tree. (b) Detailed illustration of the tree decoder’s iterative process, showing how subtree pairs are evaluated and selected. (c) Mechanism for estimating parent node representation through topology-aware fusion of local and global information. (d) Example of iterative phylogenetic tree construction using NeuralNJ, demonstrated in three stages (Steps 1-7, 8-14, and 15-20).

1. **Sequence encoder:** Sequence encoder embeds each input sequence into a high-dimension vector to capture its essential characteristics. For this aim, we implement sequence encoder using the MSA-transformer architecture (Rao et al., 2021). The advantage of this architecture is that it alternately computes attention along both species and sequence dimensions of the input MSA, and thus generates site-aware and species-aware representations (see Supplementary Algorithm 2 for further details). These representations, rather than the original genome sequences, are explored by the subsequent tree decoder for constructing phylogenetic trees.
2. **Tree decoder:** Tree decoder aims to decode a phylogenetic tree from the representations of species calculated by sequence encoder. Tree decoder starts from an initial subtree set that consists of each species as a degenerated tree with only one node, and then iteratively selects and joins two subtrees until acquiring a complete phylogenetic tree (Fig. 1b). To select the most plausible subtrees for joining, we enumerate all possible subtree pairs, estimate the embedding of the parent node of each subtree pair, and exploit this embedding to calculate the priority score that the two subtrees of this pair, if joined together, appear in the groundtruth phylogenetic tree.

It should be noted that, when estimating the embedding of the parent node of two subtrees, we consider not only the two subtrees themselves but also all other subtrees using a topology-aware gated network (Fig. 1c). This design is motivated by the key insight that a newly-added parent node (say *h*_*x*_ in Fig. 1c) serves as a topological bridge between two subtrees (*h*_*l*_ and *h*_*r*_) and other subtrees that might be merged in future steps (*h*_*rest*_). To accurately capture this bridging role, NeuralNJ employs a dual-level fusion strategy: local topology is encoded through gated left-right subtree fusion, while global ancestry information is incorporated through attention-based aggregation of remaining subtrees (see further details in Supplementary Algorithm 3).

We have also implemented three variants of NeuralNJ, including: *i*) NeuralNJ that greedily selects the subtree pair with the highest score, *ii*) NeuralNJ-MC that samples from all subtree pairs according to their scores, and *iii*) NeuralNJ-RL that uses reinforcement learning strategy with likelihood as reward rather than supervised learning. The latter two variants usually generate multiple complete trees, from which we select the highest likelihood one. Here, the likelihood of a complete phylogenetic tree is calculated using Felsenstein’s pruning algorithm (Felsenstein, 1981), which efficiently computes the likelihood by a post-order traversal of the tree.

We trained NeuralNJ on simulated data that were generated using a similar approach as Phyloformer. NeuralNJ works in an end-to-end manner, in which the training loss (the difference between a predicted tree and its ground-truth counterpart) is propagated back to the first layer. This way, both sequence encoder and tree decoder can be improved during the training pro-cess, thus facilitating learning and promoting phylogenetic inference.

NeuralNJ, once trained, can be executed directly to infer phylogenetic tree for a collection of genome sequences. We use a dataset containing 20 taxa as an example to demonstrate the phylogenetic inference process. As shown in Figure 1d, taxon 10 and 5 were selected to join together, and the joined tree was further selected to join with taxon 16. At step 8-14, the subtree consisting of taxon 10, 5, and 16 was joined with taxon 3 and 11. At step 15-20, the subtree at the bottom was joined with the subtrees on top, thus forming a complete phylogenetic tree.

### NeuralNJ achieves high accuracy on simulated data

We first evaluated NeuralNJ on simulated data that were generated using the well-established evolutionary model GTR+I+G (Tavaré, 1986; Yang, 1994). Each dataset consists of: *i*) an evolutionary tree with a random topology and branch length sampled according to an exponential distribution, *ii*) a number of taxa with sequences generated according to the evolutionary tree (see Methods for details). The evolutionary trees were used as ground-truth in subsequent evaluation procedure.

We constructed a simulated training dataset of 50-taxon phylogenetic tree-MSA pairs. For each of 29 sequence lengths (128–1024 nt in 32-nt steps), 1000 distinct tree topologies with corresponding MSAs were generated, totaling 29,000 training samples. Generally, shorter sequences represent more challenging tasks as they provide less information relative to longer sequences. Similarly, we constructed a test dataset comprising 640 samples. To evaluate the generalization ability of NeuralNJ, we set the test dataset with diverse taxa number (20, 50, 100 taxa) and sequence lengths (256, 512 nt). It should be noted that the training and test datasets were independently generated to avoid potential overlap.

We compared NeuralNJ with two state-of-the-art approaches: RAxML-NG, a well-established maximum likelihood-based solver, and Phyloformer, a machine learning-based approach. We evaluated the constructed phylogenetic trees using the normalized Robinson-Foulds (RF) distance (Robinson and Foulds, 1981), which quantifies the topological difference between a predicted tree and its ground-truth counterpart. Figure 2 presents the topological accuracy of NeuralNJ, Phyloformer, and RAxML-NG across data configurations: the left panel shows performance with fixed sequence length (1024 nt), while the right panel shows performance with fixed taxa number (100 taxa). As shown in the left panel, RAxML-NG (blue bar) and Phyloformer (purple bar) achieve mean normalized RF distances of 0.108 and 0.116, respectively, on the dataset with 20 taxa, while NeuralNJ (red bar) and its variants, NeuralNJ-MC (orange bar) and NeuralNJ-RL (yellow bar), achieve distances of 0.114, 0.089, and 0.087, respectively. The right panel shows that for sequences of length 256 nt, RAxML-NG achieves a normalized RF distance of 0.316, Phyloformer achieves 0.300, while NeuralNJ, NeuralNJ-MC, and NeuralNJ-RL achieve distances of 0.269, 0.268, and 0.262, respectively. These results demonstrate the higher accuracy of our NeuralNJ approach over RAxML-NG and Phyloformer. The performance advantage of NeuralNJ becomes more pronounced in challenging scenarios with larger numbers of taxa (e.g., 100) or shorter sequence lengths (e.g., 256 nt).

**Figure 2.**
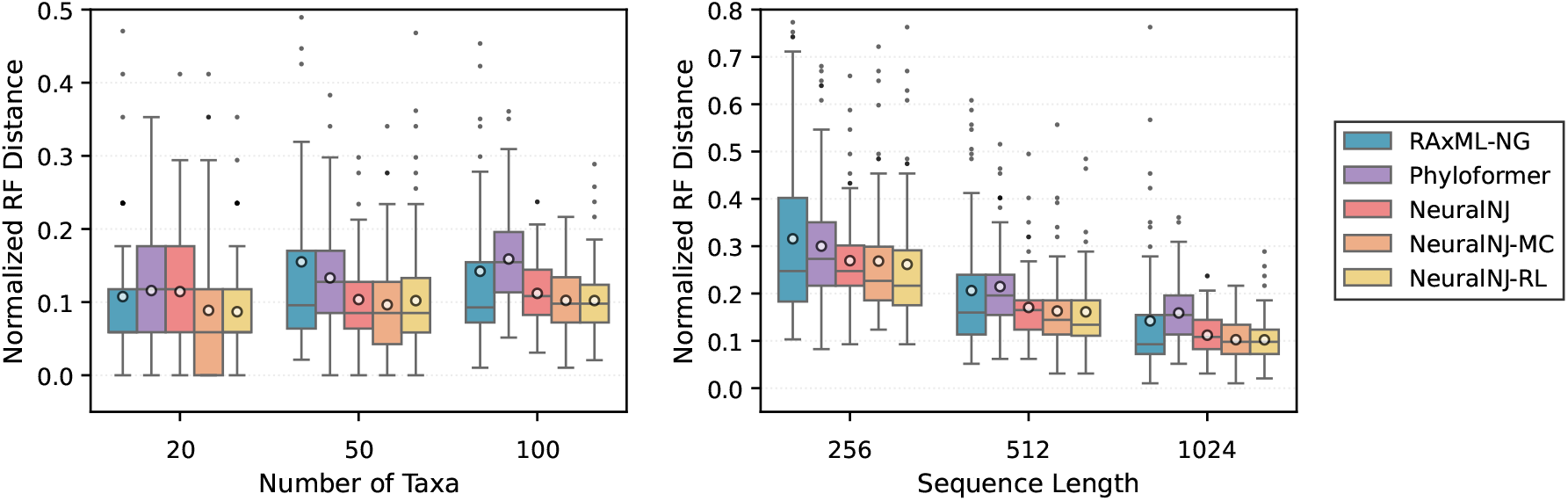
Performance comparison of phylogenetic inference methods on the simulated test dataset under the GTR+I+G evolutionary model. Normalized RF distances between predicted and ground-truth simulated trees across varying numbers of taxa (left) and sequence lengths (right). Lower RF distances indicate higher topological accuracy. Box plots show the median (central line), quartiles (box), 1.5× interquartile range (whiskers), outliers (individual points), and the mean value (circle) for each group.

In summary, these results highlight the superior performance of NeuralNJ in topological accuracy, particularly for the challenging cases with larger numbers of taxa or shorter sequence lengths. Further experiments in the Supplementary Materials demonstrate NeuralNJ’s robust performance across diverse simulation scenarios, including datasets with longer sequences and different evolutionary parameters. These evaluations consistently validate NeuralNJ’s advantages in phylogenetic tree reconstruction.

### NeuralNJ achieves high efficiency on empirical data

To validate our method’s accuracy and efficiency on real data, we evaluated it using four empirical phylogenomic DNA datasets. These datasets represent diverse taxonomic groups (Zhou et al., 2017): WickD3b with 103 land plant species, JarvD5a with 48 bird species, SongD1 with 37 mammal species, and TarvD7 with 36 mammal species. Each dataset consists of multiple gene alignment sequences from these species. Both NeuralNJ and Phyloformer, trained on the aforementioned GTR+I+G simulated data, were evaluated on these empirical datasets with varying sequence lengths based on hardware constraints: sequences up to 17,000 nucleotides for datasets with <50 taxa (achieving >98% coverage for JarvD5a, SongD1, and TarvD7) and up to 4,400 nucleotides for WickD3b (50% coverage). Results for NeuralNJ-RL are not included in the current empirical evaluation because this variant requires increased computational and hardware resources.

Due to the absence of true phylogenetic trees for empirical data, we followed the common practice of using widely accepted reference trees supported by established classical methods (Zou et al., 2019). Following Phyloformer’s evaluation protocol (Nesterenko et al., 2025), we used reference trees inferred by ML-based solvers on concatenated multiple sequence alignments as ground truth (Zhou et al., 2017). These reference trees may inherently favor ML-based methods, potentially making it difficult for our approach to demonstrate superior performance on such data.

We present our accuracy results in Figure 3. As expected, RAxML demonstrates stronger alignment with the reference trees. On JarvD5a, NeuralNJ achieves a mean RF distance of 0.78, approaching RAxML’s performance of 0.74. For SongD1, TarvD7, and WickD3b datasets, NeuralNJ shows competitive performance with mean RF distances of 0.45, 0.55, and 0.58, respectively, which are comparable to RAxML’s results. NeuralNJ outperforms Phyloformer on three datasets (JarvD5a, SongD1, and TarvD7), while showing slightly higher RF distances on WickD3b. NeuralNJ-MC, which incorporates likelihood-guided sampling strategies, further improves upon the performance of NeuralNJ. Notably, on the JarvD5a and WickD3b datasets, NeuralNJ-MC achieves lower RF distances than RAxML. Overall, NeuralNJ-MC attains comparable or marginally better accuracy, with a mean RF distance of approximately 0.53 versus 0.55 for RAxML, highlighting the potential effectiveness of our sampling-based approach. Detailed quantitative results for both the complete empirical datasets and the long-sequence subsets are provided in the Supplementary Materials.

**Figure 3.**
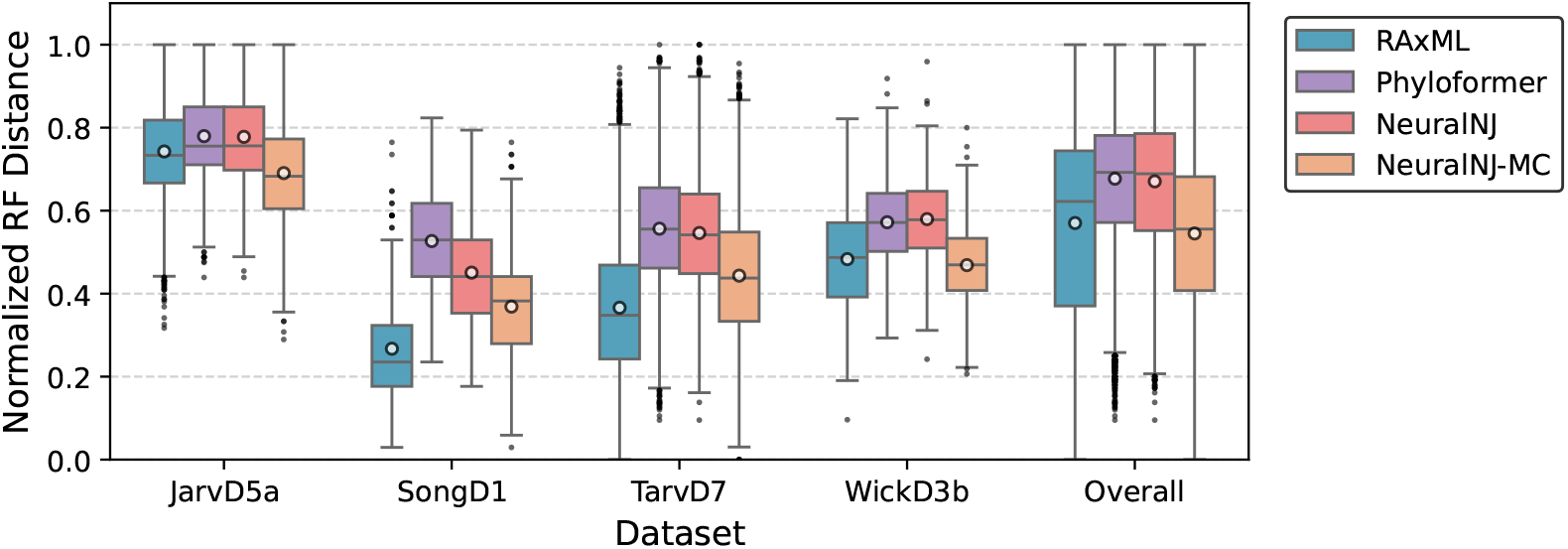
Performance evaluation on empirical phylogenomic datasets. Box plots show the distribution of normalized RF distances between inferred and reference phylogenies for different inference methods across four empirical datasets (JarvD5a, SongD1, TarvD7 and WickD3b) and their overall performance. Box plots show the median (central line), quartiles (box), 1.5× interquartile range (whiskers), outliers (individual points), and the mean value (circle) for each group, with lower RF distances indicating higher accuracy in topology reconstruction.

We present our efficiency results in Table 1. Both NeuralNJ and Phyloformer demonstrate superior inference speed, with runtime reductions of 100-200x compared to RAxML. This advantage is particularly pronounced on WickD3b, the dataset with the largest number of species, where the average inference time decreases from 184.26 seconds (RAxML) to below 1 second (NeuralNJ and Phyloformer). While NeuralNJ-MC achieve improved accuracy, this comes at the cost of increased computational time, as evidenced by their long runtimes across all datasets.

**Table 1.** Average runtime comparison (seconds) of different phylogenetic inference methods on empirical datasets.

| Dataset | RAxML * | Phyloformer | NeuralNJ | NeuralNJ-MC |
| --- | --- | --- | --- | --- |
| JarvD5a | 203.06 | 0.95 | 0.97 | 133.30 |
| SongD1 | 148.82 | 1.01 | 1.37 | 177.40 |
| TarvD7 | 99.12 | 0.74 | 0.97 | 151.34 |
| WickD3b | 184.26 | 0.95 | 0.91 | 105.26 |

\* RAxML timings were obtained from Supplementary Table S6 of the empirical data (Zhou et al., 2017), which reports runtimes on single-gene alignments.

In summary, NeuralNJ demonstrates high computational efficiency while maintaining comparable accuracy to state-of-the-art methods on empirical datasets.

### NeuralNJ perfectly balances accuracy and efficiency

Both computational efficiency and inference accuracy are critical considerations for phylogenetic inference methods, particularly given the increasing scale of modern phylogenetic analyses involving large numbers of taxa and long sequences. We evaluated the trade-off between these two factors across varying problem scales under the GTR+I+G model. Our evaluation compares NeuralNJ variants and neural baselines implemented on GPU platforms against classical solvers on CPUs (detailed hardware configurations are provided in Supplementary Table 13).

Figure 4 presents an evaluation using Pareto frontiers, which represent the optimal trade-off curve where no method can improve one metric without degrading another. The results are shown across varying numbers of taxa (Figures 4a-c) and sequence lengths (Figures 4d-f). Our analysis demonstrates that NeuralNJ consistently achieve Pareto-optimal frontiers across all experimental settings. This is particularly evident in Figure 4c, which represents a challenging scenario with 100 taxa and 1024 nt sequences, where NeuralNJ and its variants lie on the Pareto-optimal frontier. In contrast, Phyloformer falls off the frontier with a second-order runtime and an RF distance of approximately 0.16, while RAxML-NG is suboptimal with minute-scale computational time and an RF distance of approximately 0.14. While BIONJ also maintains a position on the Pareto frontier, it demonstrates substantially lower accuracy with an RF distance of approximately 0.4 under the challenging conditions of 100 taxa and 1024 nt sequences.

**Figure 4.**
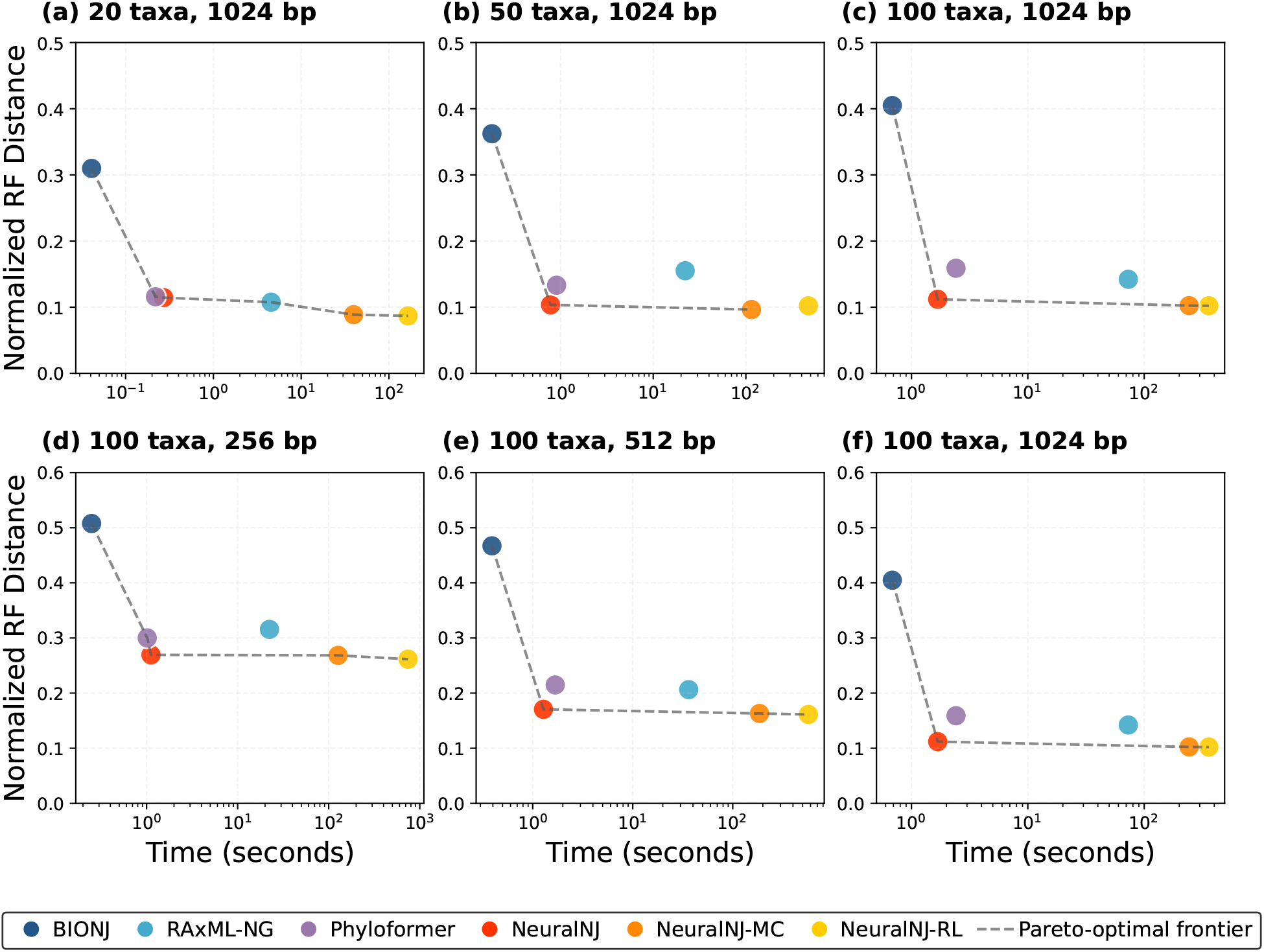
Efficiency-accuracy trade-offs of phylogenetic inference methods under the GTR+I+G model. (a-c) Accuracy versus runtime across taxon counts with fixed sequence length (1024 nt). (d-f) Comparison across sequence lengths with fixed taxon number (100). The x-axis shows computational time in seconds (log scale), and the y-axis shows the normalized Robinson-Foulds distance. Colored markers represent different methods: BIONJ (blue), RAxML-NG (light blue), Phyloformer (purple), NeuralNJ (red), NeuralNJ-MC (orange), and NeuralNJ-RL (yellow). The gray dashed line (Pareto frontier) represents the optimal trade-off between accuracy and computational time, where no method can improve one metric without compromising the other.

The scalability analysis reveals that NeuralNJ maintains its Pareto-optimal performance across increasing problem scales. Specifically, as the number of taxa increases from 20 to 100 (Figures 4a-c), NeuralNJ and its variants consistently define the efficiency-accuracy frontier, while other methods deviate from optimal performance. Similarly, across increasing se-quence lengths from 256 to 1024 nt (Figures 4d-f), NeuralNJ demonstrate robust accuracy while maintaining their computational efficiency advantage. Detailed performance analyses across different problem scales are provided in Supplementary Section D.5.

While NeuralNJ and its variants (NeuralNJ-MC/RL) consistently achieve Pareto-optimal performance, they occupy different strategic positions on the frontier. NeuralNJ establishes an optimal balance point, delivering significant accuracy improvements over traditional distance-based methods while maintaining computational efficiency. In contrast, NeuralNJ-MC and NeuralNJ-RL extend toward the high-accuracy region of the frontier, offering viable alternatives for applications where precision takes precedence over computational cost.

This evaluation demonstrates that NeuralNJ achieves an optimal balance between accuracy and efficiency across diverse problem scales, while its variants provide flexible options for different accuracy-efficiency trade-offs in phylogenetic inference tasks.

### The advantages of NeuralNJ: case studies

Figure 5 presents a case study using a simulated MSA data (50 taxa, 1024nt) under GTR+I+G model to visualize the reconstruction accuracy of different methods. The MSA data and corresponding true tree topology used in this case study are provided in Supplementary Section E. The figure compares the true phylogenetic tree (a) with trees inferred by five methods (b-f), where red branches indicate topological inconsistencies with the true tree - specifically, join operations that do not match any valid subtree combinations in the true topology.

**Figure 5.**
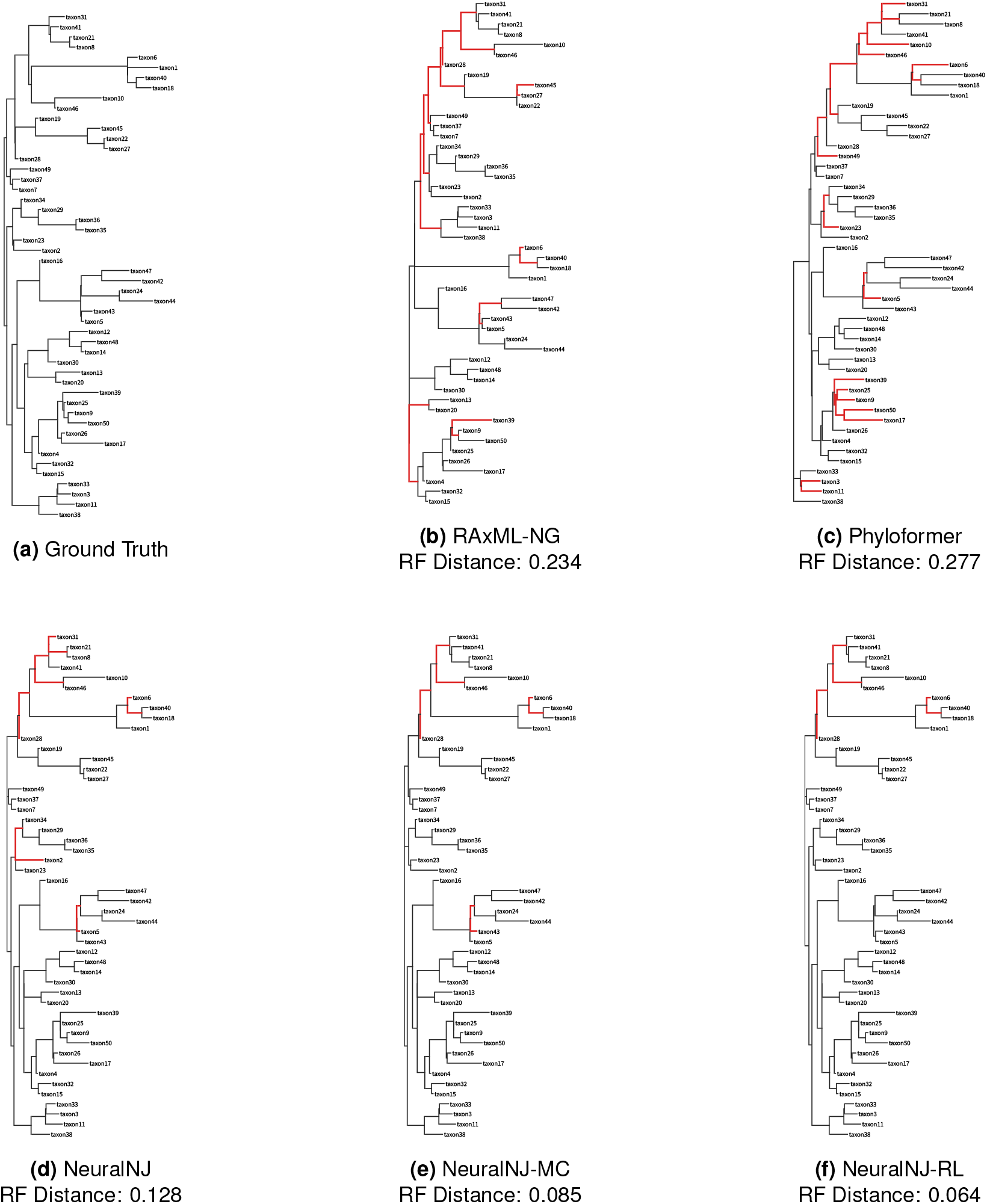
Comparison of phylogenetic tree reconstructions across different inference methods on a simulated dataset (50 taxa, 1024 nt) under the GTR+I+G model. (a) Ground truth phylogenetic tree used for simulation. (b-f) Trees inferred by RAxML-NG, Phyloformer, NeuralNJ, NeuralNJ-MC, and NeuralNJ-RL, respectively. Topological discordance with the ground truth tree is highlighted in red, while black branches indicate conserved relationships. The normalized Robinson-Foulds (RF) distances between each inferred tree and the ground truth are shown. The sequence alignment used in this analysis is provided in Supplementary Section E.

As illustrated in Figure 5, RAxML-NG erroneously clusters taxon39 with taxon9 and taxon50, while Phyloformer misplaces taxon50 by merging it with taxon17; in the reference tree, taxon50 should be grouped with taxon9. NeuralNJ and its variants correctly resolve the relationships involving taxon39 and taxon50, but some errors remain: NeuralNJ incorrectly merges taxon2 with the cluster containing taxon34, taxon29, taxon36, and taxon35. NeuralNJ-MC corrects this error, but misorders the merging of taxon43 and taxon5. NeuralNJ-RL further improves the reconstruction, achieving the lowest RF distance (0.064) among all methods. Overall, RAxML-NG and Phyloformer exhibit substantial topological errors (RF distances of 0.234 and 0.277, respectively), while the NeuralNJ variants demonstrate progressively improved reconstruction accuracy, with RF distances decreasing from 0.128 (NeuralNJ) to 0.085 (NeuralNJ-MC) and 0.064 (NeuralNJ-RL).

Figure 6 illustrates the dynamic tree construction processes of NeuralNJ and distance-based neighborjoining (NJ) methods. Using simulated multiple sequence alignment (MSA) data comprising 50 taxa and 1024nt generated under the GTR+I+G model (provided in Supplementary Section F), we examined an ideal case where NeuralNJ achieves correct topology prediction and NJ uses the true distance matrix from the reference tree.

**Figure 6.**
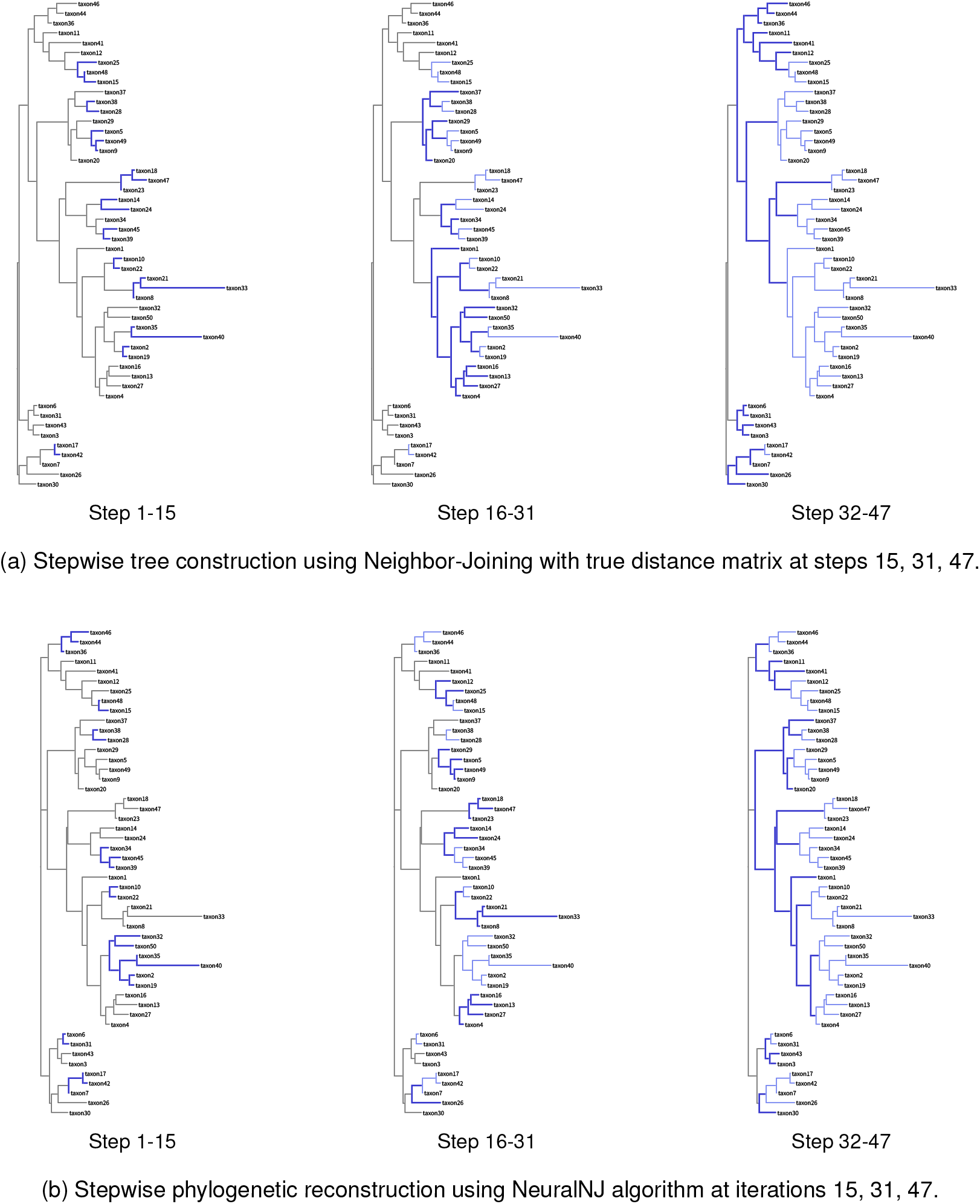
Stepwise phylogenetic tree reconstruction comparing Neighbor-Joining and NeuralNJ on simulated sequence data (50 taxa, 1024 nt) under the GTR+I+G model. (a) Tree construction process using classical Neighbor-Joining with the true evolutionary distance matrix and (b) NeuralNJ algorithm, shown at steps 15, 31, and 47. Branch colors denote merging order: black represents unmerged taxa, blue indicates recent merging events (within previous 10 steps), and light blue shows earlier merging events. Both methods converge to identical topologies (Robinson-Foulds distance = 0). The sequence alignment used in this analysis is available in Supplementary Section F.

The NJ algorithm demonstrates a region-focused construction pattern. In the initial steps (1–15), it merges deep leaf nodes such as taxon10 and taxon22, or small clusters like taxon21, taxon33, and taxon8. During steps 16–31, these small clusters are further joined with other taxa, including taxon1, taxon32-taxon19 and taxon16-taxon4, which all belong to the same major clade. This is followed by the progressive assembly of deeper internal nodes, ultimately forming the central topology of the tree. The Q-matrix calculation mechanism of NJ algorithm preferentially constructs branches that are close to each other while maintaining considerable distance from other sequences in the dataset. Although the NJ method guarantees accurate tree reconstruction when distance estimates are precise, this stepwise joining behavior may potentially lead to error accumulation in practical applications where distance estimates contain inaccuracies. Our NeuralNJ method exhibits a distinctive construction pattern where merge operations are distributed across different regions of the tree simultaneously. As shown in the Figure 6 (Steps 1-15 and 16-31), the algorithm concurrently processes multiple regions, merging taxon28 and taxon40 in the upper region, taxon34 and taxon39 in the middle, and taxon17 and taxon7 in the lower region. This distributed merging strategy differs fundamentally from NJ’s region-focused approach. This behavior emerges from NeuralNJ’s learning objective: trained on labeled trees, it learns to identify and prioritize correct pair-wise relationships globally, regardless of their regional location. Consequently, the algorithm achieves a more balanced tree construction process, potentially reducing the risk of error propagation that could occur from consecutive merges within a single region.

## Discussion

This study presents NeuralNJ, demonstrating strong performance in phylogenetic inference by achieving an optimal balance between accuracy and efficiency. The end-to-end neural architecture enables direct topology learning and efficient single-pass tree construction, eliminating error accumulation inherent in iterative optimization approaches. Experiments on simulated datasets of varying scales demonstrate that NeuralNJ is robust and particularly excels in complex scenarios and large-scale problems with 100 species. On empirical datasets, NeuralNJ consistently shows competitive accuracy while significantly reducing computational overhead. Case studies further validate that our end-to-end distributed tree construction effectively minimizes error propagation, enhancing the overall accuracy of phylogenetic reconstruction.

The exceptional performance of NeuralNJ stems from three key architectural innovations: an end-to-end neural framework for training and tree construction, learning-to-rank strategy for join operations, and topology-aware parent node representation estimation. First, the end-to-end architecture enables direct optimization towards ground-truth tree topology reconstruction, while eliminating the time-consuming iterative topology optimization through single-pass inference. Second, learning join priorities instead of direct distance matrix estimation offers dual advantages: it simplifies the learning objective by focusing on correct pairing order rather than precise distance values, enhancing model scalability; and it allows the model to concentrate exclusively on high-priority pairs, making the learning process more focused and efficient. Third, our topology-aware parent node representation incorporates not only immediate child node features but also innovatively integrates ancestral context, with ablation studies demonstrating significant performance improvements from this contextual integration.

While NeuralNJ effectively learns to map genome sequences to tree topologies from simulated data, its performance is inherently bounded by data quality and the similarity between target problems and simulation data. We observe notable disparities in accuracy between simulated and empirical datasets, with empirical data showing consistently higher normalized RF distances. This gap likely stems from several factors in the current simulation-based training paradigm. First, the scarcity of real-world labeled trees prevents direct training on empirical data. Second, the GTR+I+G evolutionary model used in our simulations may not optimally fit all empirical datasets, despite evolutionary model selection being crucial for phylogenetic inference. Traditional phylogenetic inference typically involves optimal model selection, an aspect that remains unexplored in the context of neural networks and simulated data. Additionally, our current approach of using uniform tree topology distributions and exponential branch length distributions may not fully capture real-world evolutionary patterns, suggesting the need for more sophisticated simulation strategies aligned with empirical data distributions.

Despite the efficient computation of neural network methods for phylogenetic inference, their broader application remains constrained by hardware limitations. Due to GPU memory constraints, current neural network methods, such as NeuralNJ and Phyloformer, are typically limited to processing sequences shorter than 5,000 nucleotides for problems involving 100 taxa, and sequences under 17,000 nucleotides for datasets with 50 taxa. These technical limitations significantly restrict the application of neural approaches to large-scale phylogenomic datasets, which often contain concatenated alignments spanning hundreds of thousands to millions of sites. Future advancements in this field will necessitate the development of memory-efficient architectures capable of handling substantially longer sequence alignments while maintaining computational tractability.

NeuralNJ’s flexible framework integrates both likelihood-free inference and likelihood-guided optimization within a unified approach. NeuralNJ directly infers phylogenetic tree topologies from sequence data, effectively circumventing the computationally expensive likelihood calculations that constrain maximum likelihood methods when analyzing complex evolutionary models Nesterenko et al. (2025). However, the base NeuralNJ model, trained on trees sampled from predefined evolutionary distributions, is fundamentally limited by its reliance on the training prior, which constrains its generalization to out-of-distribution data and may lead to suboptimal solutions for specific empirical cases. To address this limitation, NeuralNJ-MC and NeuralNJ-RL variants incorporate likelihood-based tree selection and reinforcement learning optimization, enabling the model to adapt its search beyond the constraints of the training prior and improve performance on individual instances. Experimental results indicate that calibrated likelihood guidance generally enables models to explore broader tree space and can optimize both likelihood and topological accuracy simultaneously. However, representative cases presented in the Supplementary Material show that likelihood optimization does not always guarantee improved RF distance, and the relationship between likelihood and topology remains complex and case-dependent. These results underscore the complexity of the relationship between likelihood and topology, indicating promising directions for future research.

While NeuralNJ-MC and NeuralNJ-RL demonstrate superior accuracy compared to the base NeuralNJ model, this improvement comes at the cost of intensive likelihood calculations and branch length optimizations. Notably, the computational demands of NeuralNJ-RL’s fine-tuning process are particularly significant, requiring substantial GPU resources for each case, which emphasizes the necessity for more efficient search strategies. Future research directions could productively explore neural approaches for direct branch length prediction and efficient likelihood estimation, while simultaneously investigating methods to reduce the impact of phylogenetic prior biases in the training data.

Our work on NeuralNJ, together with its variants and recent developments in deep learning approaches for phylogenetic inference, represents a significant step towards efficient and accurate evolutionary tree reconstruction. The demonstrated balance between computational efficiency and accuracy, particularly in complex evolutionary scenarios, will contribute to better understanding of evolutionary relationships across diverse species. Future developments in simulation data quality and neural network architectures will further enhance the applicability of deep learning methods in phylogenetic analysis, potentially revolutionizing our approach to studying evolutionary history.

## Materials and methods

### Data and implementation

#### Simulated phylogenetic data

We generated simulated phylogenetic data using a three-stage approach informed by empirical observations. First, tree topologies were generated using uniform random binary trees via Biopython (Cock et al., 2009; Talevich et al., 2012). Next, branch lengths were sampled from exponential distributions with rate parameters *λ* (Rannala and Yang, 1996; Stadler, 2011) derived from real phylogenetic datasets (Azouri et al., 2021). Based on the analysis of these empirical data, *λ* values were drawn uniformly from [log 2, log 5] in log space. DNA sequences were simulated with IQ-TREE2’s Alisim (Ly-Trong et al., 2022, 2023) under the GTR+I+G model (Tavaré, 1986). For each alignment, substitution rates, base frequencies, gamma shape, and invariant site proportions were sampled from empirical distributions (Naser-Khdour et al., 2021). To reproduce realistic gap patterns, deletion rates were varied within [0, 0.01, 0.02, 0.03, 0.04], and rates below 0.04 were chosen to keep simulated gap ratios below 0.3, consistent with empirical observations (10–30%)(Zhou et al., 2017).

The training dataset consists of 29,000 unique tree-MSA pairs, with each phylogenetic tree simulated independently and paired with exactly one corresponding MSA. All training samples contain 50 taxa, while sequence lengths vary from 128 to 1024 nucleotides in increments of 32 (resulting in 29 different sequence lengths, with 1000 unique tree-MSA pairs per length). For the test dataset, we similarly generated unique tree-MSA pairs (each consisting of one independently simulated phylogenetic tree and its corresponding MSA) according to the following design: 128 pairs for each combination of sequence length 1024 and taxa counts of 20, 50, and 100; additionally, for the 100-taxa case, we included 128 pairs each for sequence lengths of 256 and 512 nucleotides.

#### Evolutionary models

We employed the GTR+I+G model for our main analyses, with all model parameters (substitution rates, base frequencies, gamma shape, and invariant site proportion) sampled from empirical distributions (Naser-Khdour et al., 2021). This approach allows for varying substitution rates among nucleotides, rate heterogeneity across sites (+G), and invariable sites (+I), making it particularly suitable for real-world phylogenetic analyses (Sumner et al., 2012; Arenas, 2015). Additional analyses incorporating using the sim-pler Jukes-Cantor (JC) model (Jukes et al., 1969) are presented in the Supplementary Materials.

#### Empirical datasets

We evaluated NeuralNJ on empirical phylogenomic data from Zhou et al. (Zhou et al., 2017), which included four distinct DNA collections: WickD3b (land plants), JarvD5a (birds), SongD1 (mammals), and TarvD7 (mammals). For datasets with fewer than 50 taxa (JarvD5a, SongD1, TarvD7), we included sequences up to 17,000 nucleotides; for WickD3b (up to 100 taxa), we processed sequences up to 4,400 nucleotides. This approach achieved thorough dataset coverage: SongD1 (100%), JarvD5a (98%), TarvD7 (99.8%), and WickD3b (50%, limited by available reference trees). For each dataset, we used the reference tree topology inferred by maximum likelihood estimation on concatenated genome sequences (Nesterenko et al., 2025).

#### Baselines for comparison

We selected a range of baseline methods for phylogenetic inference, including the heuristic distance-based algorithm BIONJ (Gascuel, 1997), the well-established maximum likelihood phylogenetic solver RAxML-NG (Kozlov et al., 2019), and the neural network-based inference algorithm Phyloformer (Nesterenko et al., 2025). BIONJ represents a commonly used heuristic approach for initial tree generation in evolutionary solvers. RAxML-NG represents a highly optimized, mature phylogenetic solver. Phyloformer, a recent development, exemplifies state-of-the-art neural network approaches in this field. By comparing our proposed method with these established techniques, we aimed to comprehensively evaluate its performance and effectiveness in phylogenetic inference. It is worth noting that both NeuralNJ and Phyloformer were trained on identical training data and evaluated using a consistent early stopping mechanism (training terminates when validation performance shows no improvement for five consecutive validation checks) to ensure fair comparison.

#### Evaluation metric

We primarily evaluated NeuralNJ using the Normalized Robinson-Foulds (RF) distance (Robinson and Foulds, 1981), which measures the structural differences between inferred and reference trees. This metric directly quantifies the topological accuracy of phylogenetic inference methods by comparing the shared splits between two trees, with values ranging from 0 (identical topologies) to 1 (completely different topologies). Additional analyses, including likelihood-based evaluations under different branch length optimization strategies, are presented in the Supplementary Materials.

### Model design of NeuralNJ

The proposed NeuralNJ framework comprises three essential neural modules for phylogenetic tree construction: (1) a sequence encoder (ℳ _enc_) that transforms MSA input into learned sequence representations, and a tree decoder consisting of (2) a parent node representation estimator (ℳ_est_) that conducts topology-aware representation estimation of potential ancestral nodes for candidate subtree pairs, and (3) a subtree evaluator ( ℳ_eval_) that determines joining priorities based on these ancestral representations. The architectural details of these modules are elaborated in the following sections, while their complete mathematical formulations are presented in Supplementary Section C.

#### (1) Sequence encoder ℳ_Enc_

The sequence encoder ℳ_Enc_ in NeuralNJ adopts the axial attention architecture from MSA-Transformer (Rao et al., 2021), which effectively captures evolutionary relationships between species while accommodating variable sequence lengths and quantities.

The encoding process consists of three main steps. First, for an input MSA 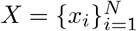 containing *N* se-quences, each sequence *x*_*i*_ of length *L* is embedded into a high-dimensional space through a multilayer perceptron (MLP) with a GELU-activated hidden layer, yielding embeddings 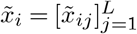.

Second, these embeddings undergo transformation through a cascade of axial attention layers (Ho et al., 2019), following the MSA-Transformer architecture (Rao et al., 2021). Each layer implements two complementary attention mechanisms: *i*) column attention, which computes position-wise correlations between species embeddings, and *ii*) tied row attention, which employs a shared attention map across species to capture positional correlations within individual sequences. Both mechanisms operate with a memory complexity of *O*(*LN* ^2^ + *L*^2^).

Finally, the encoder produces 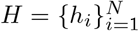 for the *N* species, where each species’ representation *h*_*i*_ is a composite of its sequence embedding representations after the attention transformations, i.e., 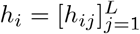. This encoding process facilitates the transfer of information across species sequences, effectively capturing inter-species evolutionary information.

#### (2) Parent node representation estimator ℳ_Est_

As illustrated in Figure 1c, the parent node representation estimator ℳ_Est_ computes the parent node representation *h*_*x*_ for two subtrees with representations *h*_*l*_ and *h*_*r*_ that are to be joined. In phylogenetic tree construction, the parent node serves as a bridge between its immediate descendants and more distant ancestral nodes. This dual role imposes two key requirements: First, *h*_*x*_ must maintain strong correlations with both *h*_*l*_ and *h*_*r*_, as it represents their immediate common ancestor. Second, *h*_*x*_ should also capture relationships with potential ancestral and sibling nodes, whose information is embedded within the broader subtree set excluding *h*_*l*_ and *h*_*r*_.

To achieve topology-aware parent node representation estimation that satisfies these requirements, ℳ_Est_ computes *h*_*x*_ through a two-stage process through hierarchical aggregation:

##### Stage 1: immediate descendant integration

A lightweight gating network performs site-by-site aggregation of the subtree representations 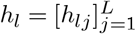 and 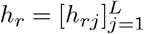 to generate an intermediate representation 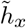:

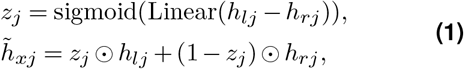

where *j* denotes the site index, and the gating mechanism adaptively weights the contribution of each subtree’s features.

##### Stage 2: ancestral context integration

The final representation *h*_*x*_ is computed by integrating 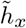 with a context-aware representation *h*_rest_ through another sitespecific gating network:

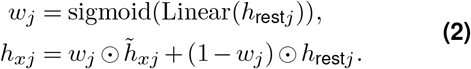

The context-aware representation *h*_rest_ encapsulates information from potential ancestral and sibling nodes by aggregating the complementary subtree set (excluding *h*_*l*_ and *h*_*r*_). This aggregation is implemented through a cross-attention mechanism (Vaswani et al., 2017), using 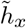 as the query and the remaining subtree representa-tions {*h*_*k*_, *k* ≠ *l, r*} as keys:

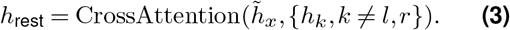

Notably, while the gating networks operate site-wise to preserve fine-grained sequence features, the crossattention mechanism processes complete sequence representations to maintain comprehensive evolutionary information in *h*_rest_. The detailed computational procedure is provided in Algorithm 3 in Supplementary Section C.

Through this hierarchical two-stage process, the representation aggregator efficiently computes the parent node representation 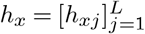. The combination of site-wise gating operations and cross-attention mechanism ensures that *h*_*x*_ effectively captures both local relationships with direct descendants and broader evolutionary context from potential ancestors and siblings. Importantly, the adaptive gating design maintains gradient stability during backpropagation, allowing robust optimization even for deep phylogenetic trees with large species sets. As demonstrated in Supplementary Figure S18, this two-stage aggregation process with attention-based ancestral context integration achieves superior convergence compared to simpler alternatives like child-only aggregation or sum-based integration. This topology-aware representation lays the foundation for subsequent priority prediction and tree construction.

#### (3) Subtree Evaluator ℳ_Eval_

The subtree evaluator maps the aggregated node representation *h*_*x*_ to a scalar priority score 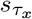 for the newly-joined subtree *τ*_*x*_. Specifically, a MLP first transforms each site-specific feature of 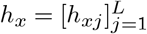 into a scalar value, and these values are then averaged across all sites to produce the final priority score. This site-wise averaging strategy not only ensures computational efficiency but also provides balanced consideration of evolutionary signals from all sites.

### Subtree selection strategies

NeuralNJ’s flexible framework supports multiple strategies for selecting subtree pairs based on the predicted priority scores 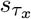. NeuralNJ employs a direct greedy selection strategy, while NeuralNJ-MC and NeuralNJ-RL utilize probabilistic selection mechanisms to enable likelihood-guided search.

The greedy strategy, implemented in NeuralNJ, directly selects the newly-joined subtree with the highest priority score:

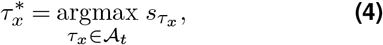

where 𝒜_*t*_ represents the set of all candidate newly-joined subtree at step *t*.

For enhanced topology exploration in NeuralNJ-MC and NeuralNJ-RL, we implement probabilistic selection strategies. These approaches convert priority scores into sampling probabilities using a temperature-scaled softmax function. For notational convenience, we express this probability distribution as:

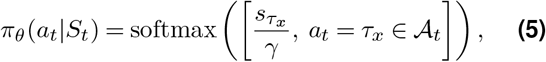

where *S*_*t*_ denotes the subtree set at step *t, γ* is the sampling temperature that controls exploration breadth. This probabilistic framework serves as the foundation for likelihood-guided search through Monte-Carlo sampling and reinforcement learning, respectively.

### Supervised training and reinforced search

#### Supervised training

To train the NeuralNJ algorithm supervisedly using the existing simulated MSA phylogenetic tree dataset, we design a training algorithm based on NeuralNJ’s bottom-up tree construction process. Given an MSA and its corresponding ground-truth tree *τ*_gt_, we reconstruct *τ*_gt_ with NeuralNJ and optimize the priority scores at each construction step.

At the *t*-th step, we identify two sets of candidate newly-joined subtrees: those that are part of *τ*_gt_ and those that are not. To ensure accurate reconstruction, candidates belonging to *τ*_gt_ should receive higher priority scores. We formalize this intuition using a binary margin-based pairwise ranking loss, which adopts the hinge loss formulation to enforce a margin between positive (ground-truth) and negative subtree candidates (Burges et al., 2005):

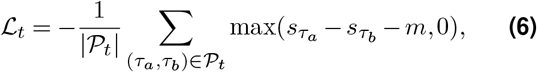

where 𝒫_*t*_ denotes all pairs (*τ*_*a*_, *τ*_*b*_) with *τ*_*a*_ ∈ 𝒜_*t*_ being a correct subtree (part of *τ*_gt_) and *τ*_*b*_ ∈ 𝒜_*t*_ being an incorrect one at step *t, s*_*τ*_ represents the priority score for subtree *τ*, and *m* is the margin hyperparameter.

This ranking loss is theoretically motivated by the RF distance, which quantifies topological differences between two phylogenetic trees by counting the number of unique splits present in one tree but not in the other (Robinson and Foulds, 1981). By encouraging higher priority scores for correct splits while penalizing incompatible ones at each step, our loss function effectively guides the tree construction process to maximize the number of shared splits between the reconstructed tree and the ground truth tree, thereby minimizing their RF distance.

This loss function design directly supports our greedy selection strategy. By requiring all correct sub-trees to have scores at least m higher than all incorrect subtrees, when this margin condition is satisfied across all possible pairs, the highest-scoring subtree will necessarily be from the set of correct subtrees. This formulation allows the model to focus on difficult cases where the margin is insufficient while ignoring already well-predicted pairs, improving training stability and effectiveness. The margin-based approach effectively teaches the neural network to prioritize correct joining operations during inference, making our greedy selection strategy both efficient and accurate.

Notably, among the candidate newly-joined subtrees 𝒜_*t*_, incorrect candidates typically constitute a large majority. We therefore adopt a selective penalization strategy, focusing on incorrect candidates with higher priority scores. This approach serves two purposes: it enhances the model’s accuracy in identifying high-priority subtrees, thereby improving overall tree construction accuracy, while also reducing the cardinality of 𝒫_*t*_, leading to optimized computational overhead and increased training efficiency. The detailed selection techniques are elaborated in the Implementation Details section of the Supplementary Material.

The overall supervised learning objective is obtained by averaging the ranking losses across both the batch samples and tree construction steps:

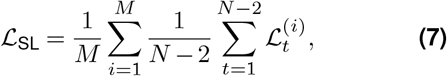

where *M* denotes the batch size, *N* is the number of species, and *N* − 2 represents the total number of joining operations required to construct a complete binary tree.

#### Reinforced search

NeuralNJ-RL leverages reinforcement learning to enhance its tree construction strategy for specific phylogenetic inference tasks, naturally aligning with the sequential decision-making nature of tree construction. While our supervised learning phase minimizes the RF distance to simulated ground truth trees, this approach may have limited generalization capability when encountering evolutionary patterns that deviate from the simulation assumptions. To enhance model adaptability to the specific case we employ phylogenetic likelihood log *P* (*X*| *τ* ) (Felsenstein, 1981) as the reward signal, directly optimizing this well-established measure that captures the fit between the tree topology and the specific observed sequence data. By adopting the REINFORCE policy gradient algorithm (Williams, 1992) with the probabilistic selection mechanism defined in Equation 5, it effectively balances exploration of diverse tree topologies with exploitation of promising construction paths. Its learning objective is defined as:

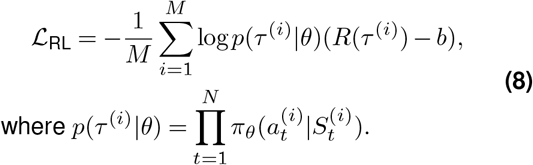

Here, *M* denotes the batch size, *p*(*τ* ^(*i*)^ | *θ*) represents the likelihood of constructing the *i*-th sampled tree under parameters *θ. R*(*τ* ^(*i*)^) = log *P* (*X* | *τ* ^(*i*)^) is the reward, computed as the phylogenetic likelihood of the final topology, and *b* is a baseline reward estimated from the previous epoch to reduce gradient variance. By maximizing this objective, the model learns to assign higher probabilities to construction paths that lead to trees with greater phylogenetic likelihood. This likelihood-guided learning enables the model to capture the long-term consequences of its joining decisions in the vast space of possible phylogenies.

## Acknowledgements

XZ, SD, CY, JZ and DB acknowledge the supports from the National Key Research and Development Program of China (2024YFC3405500) and the National Natural Science Foundation of China (32271297, 82130055). The numerical calculations in this study were supported by ICT Computer-X center. We utilized AI tools to assist in grammar correction and translation during the preparation of this manuscript.

## Data Availability

All data, code, and analysis files associated with this study are freely available at https://github.com/ZhangXinru99/NeuralNJ.

## Appendix

### Technical primer for NeuralNJ

#### Transformer architectures

The transformer is a neural network design that excels at processing sequential data (like DNA/protein sequences) by letting all elements communicate directly through attention mechanisms including self-attention and cross-attention. Unlike traditional neural networks such as Recurrent Neural Networks (Schuster and Paliwal, 1997) that process sequences word-by-word, or Convolutional Neural Networks (Gu et al., 2018) that analyze local patterns through fixed window sizes, transformers analyze all positions simultaneously through information exchange. The core architecture contains stacked processing units that progressively refine sequence understanding through two complementary components: 1) An encoder that integrates contextual information across the entire sequence, and 2) A decoder that generates predictions based on these learned patterns. This parallel processing enables efficient modeling of long-range evolutionary dependencies that span entire genomes or protein families. The power of this architecture is exemplified by breakthroughs like AlphaFold2 (Jumper et al., 2021), where transformer-based processing of multiple sequence alignments (MSAs) enables accurate protein structure prediction by capturing co-evolutionary patterns across distant sequence positions.

#### Cross-attention mechanism

Cross-attention facilitates adaptive information integration between biological entities through query-key-value interactions. The mathematical formulation is:

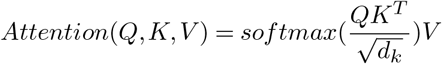

with distinct biological interpretations: Queries (Q) originate from the target representation (e.g., ancestral nodes being estimated in NeuralNJ), and Keys (K) and Values (V) derive from the source representation (e.g., sibling subtrees in NeuralNJ).

#### Self-attention mechanism

The self-attention mechanism computes contextual relationships within a single biological sequence through learnable interactions between all position pairs. It calculates attention using the same mathematical foundation as crossattention, but with Q, K, and V all derived from the same input sequence. This allows each nucleotide/amino acid position to dynamically weight its evolutionary dependencies on other sites.

#### Axial-attention mechanism

For multiple biological sequences, we often use axial-attention, a modified self-attention that separately handle row-wise (within-species) and column-wise (across-species) relationships in sequence alignments, imitating how evolutionary patterns form through both vertical inheritance and horizontal constraints. The axial-attention approach has demonstrated success in key biological applications including protein contact prediction, mutation effect forecasting, and evolutionary coupling analysis (Rao et al., 2021), making it particularly suitable for phylogenetic tasks requiring simultaneous analysis of within-species variation and cross-species conservation patterns.

#### Gated mechanism

Gating is a neural network technique that dynamically regulates information flow using learnable filters. These adaptive gates employ mathematical functions (like sigmoid) to compute precise blending ratios (0-1) between different inputs at each sequence position to preserve critical features while suppressing noise. In NeuralNJ, this enables selective integration of evolutionary signals during ancestral node estimation, where gates automatically determine which descendant mutations to retain and which ancestral patterns to emphasize. Similar gating strategies have proven essential in cutting-edge biological models like AlphaFold2, where they help balance local structural constraints with global folding patterns during protein structure prediction.

#### Multi-layer perceptron (MLP)

The MLP operates as a universal function approximator through hierarchical nonlinear transformations. Each layer performs affine projection *h*^(*l*)^ = *W* ^(*l*)^*x*^(*l*−1)^ + *b*^(*l*)^ followed by ReLU activation max(0, *h*^(*l*)^), progressively distilling high-dimensional embeddings into compact evolutionary signals.

#### NeuralNJ Architecture for Phylogenetic Inference

NeuralNJ employs a transformer-based encoder-decoder architecture specifically designed for phylogenetic tree construction. The encoder processes multiple sequence alignments (MSAs) using axial-attention mechanisms which can independently analyze vertical relationships (comparing homologous sites across species columns) and horizontal patterns (examining sequence context within individual rows). This dual-axis approach generates evolutionary embeddings that simultaneously encode cross-species conservation and lineage-specific variations.

The decoder translates these representations into tree-building decisions through a hierarchical process. Its core component, the parent estimator, operates in two phases: (1) immediate descendant integration through a lightweight gating network that combines direct child node features, and (2) sibling node aggregation using cross-attention mechanisms and gating network to integrate complementary subtree representations (excluding direct descendants). This two-stage aggregation with ancestral context integration achieves superior topological reconstruction while maintaining biologically plausible evolutionary relationships.

The subtree evaluator transforms the aggregated node representation into scalar priority scores for new subtree configurations. Specifically, a multi-layer perceptron processes the parent node’s estimated representation to generate final priority scores, which guide the selection of optimal subtree pairs during the treebuilding process.

## Bibliography

M. Arenas. Trends in substitution models of molecular evolution. Frontiers in Genetics, 6: 319, 2015.

D. Azouri, S. Abadi, Y. Mansour, I. Mayrose, and T. Pupko. Harnessing machine learning to guide phylogenetic-tree search algorithms. Nature Communications, 12(1): 1983, 2021.

D. Azouri, O. Granit, M. Alburquerque, Y. Mansour, T. Pupko, and I. Mayrose. The Tree Reconstruction Game: Phylogenetic Reconstruction Using Reinforcement Learning. Molecular Biology and Evolution, 41(6):msae105, 06 2024. doi: 10.1093/molbev/msae105. URL https://doi.org/10.1093/molbev/msae105.

M. Bordewich and C. Semple. On the Computational Complexity of the Rooted Subtree Prune and Regraft Distance. Annals of Combinatorics, 8(4):409–423, Jan 2005. doi: 10.1007/s00026-004-0229-z. URL https://doi.org/10.1007/s00026-004-0229-z.

R. Bouckaert, J. Heled, D. Kühnert, T. Vaughan, C.-H. Wu, D. Xie, M. A. Suchard, A. Rambaut, and A. J. Drummond. BEAST 2: A Software Platform for Bayesian Evolutionary Analysis. PLOS Computational Biology, 10:1–6, 04 2014. doi: 10.1371/journal.pcbi.1003537. URL https://doi.org/10.1371/journal.pcbi.1003537.

L. Brocchieri. Phylogenetic Inferences from Molecular Sequences: Review and Critique. Theoretical Population Biology, 59(1): 27–40, 2001. doi: 10.1006/tpbi.2000.1485. URL https://www.sciencedirect.com/science/article/pii/S0040580900914850.

C. Burges, T. Shaked, E. Renshaw, A. Lazier, M. Deeds, N. Hamilton, and G. Hullender. Learning to rank using gradient descent. In Proceedings of the 22nd International Conference on Machine Learning, ICML ‘05, page 89–96, New York, NY, USA, 2005. Association for Computing Machinery. ISBN 1595931805. doi: 10.1145/1102351.1102363. URL https://doi.org/10.1145/1102351.1102363.

B. Chor and T. Tuller. Maximum likelihood of evolutionary trees: hardness and approximation. Bioinformatics, 21(suppl_1):i97–i106, 2005.

P. J. Cock, T. Antao, J. T. Chang, B. A. Chapman, C. J. Cox, A. Dalke, I. Friedberg, T. Hamelryck, F. Kauff, B. Wilczynski, et al. Biopython: freely available Python tools for computational molecular biology and bioinformatics. Bioinformatics, 25(11): 1422, 2009.

R. Desper and O. Gascuel. Fast and Accurate Phylogeny Reconstruction Algorithms Based on the Minimum-Evolution Principle. In R. Guigó and D. Gusfield, editors, Algorithms in Bioinformatics, Berlin, Heidelberg, 2002. Springer Berlin Heidelberg. ISBN 978-3-540-45784-8.

J. S. Farris. Methods for Computing Wagner Trees. Systematic Biology, 19(1):83–92, 03 1970. doi: 10.1093/sysbio/19.1.83. URL https://doi.org/10.1093/sysbio/19.1.83.

J. Felsenstein. The Number of Evolutionary Trees. Systematic Biology, 27(1):27–33, 03 1978. doi: 10.2307/2412810. URL https://doi.org/10.2307/2412810.

J. Felsenstein. Evolutionary trees from DNA sequences: A maximum likelihood approach. Journal of Molecular Evolution, 17(6):368–376, Nov 1981. doi: 10.1007/BF01734359. URL https://doi.org/10.1007/BF01734359.

J. Felsenstein. Inferring phylogenies. In Inferring phylogenies, pages 664–664. 2004.

W. M. Fitch and E. Margoliash. Construction of phylogenetic trees: a method based on mutation distances as estimated from cytochrome c sequences is of general applicability. Science, 155(3760): 279–284, 1967.

O. Gascuel. BIONJ: an improved version of the NJ algorithm based on a simple model of sequence data. Molecular Biology and Evolution, 14(7): 685–695, 1997.

M. Gómez-Corrales and C. Prada. Cryptic lineages respond differently to coral bleaching. Molecular Ecology, 29(22): 4265–4273, 2020.

J. Gu, Z. Wang, J. Kuen, L. Ma, A. Shahroudy, B. Shuai, T. Liu, X. Wang, G. Wang, J. Cai, et al. Recent advances in convolutional neural networks. Pattern recognition, 77: 354–377, 2018.

S. Guindon, J.-F. Dufayard, V. Lefort, M. Anisimova, W. Hordijk, and O. Gascuel. New Algorithms and Methods to Estimate Maximum-Likelihood Phylogenies: Assessing the Performance of PhyML 3.0. Systematic Biology, 59(3):307–321, 05 2010. doi: 10.1093/sysbio/syq010. URL https://doi.org/10.1093/sysbio/syq010.

J. Ho, N. Kalchbrenner, D. Weissenborn, and T. Salimans. Axial Attention in Multidimensional Transformers. arXiv preprint arXiv:1912.12180, 2019.

S. J. Hoyt, J. M. Storer, G. A. Hartley, P. G. Grady, A. Gershman, L. G. de Lima, C. Limouse, R. Halabian, L. Wojenski, M. Rodriguez, et al. From telomere to telomere: The transcriptional and epigenetic state of human repeat elements. Science, 376(6588):eabk3112, 2022.

T. H. Jukes, C. R. Cantor, et al. Evolution of protein molecules. Mammalian Protein Metabolism, 3(24): 21–132, 1969.

J. Jumper, R. Evans, A. Pritzel, T. Green, M. Figurnov, O. Ronneberger, K. Tunyasuvunakool, R. Bates, A. Žídek, A. Potapenko, et al. Highly accurate protein structure prediction with alphafold. nature, 596(7873): 583–589, 2021.

A. M. Kozlov, D. Darriba, T. Flouri, B. Morel, and A. Stamatakis. RAxML-NG: a fast, scalable and user-friendly tool for maximum likelihood phylogenetic inference. Bioinformatics, 35 (21):4453–4455, 05 2019. doi: 10.1093/bioinformatics/btz305. URL https://doi.org/10.1093/bioinformatics/btz305.

J. Li, S. Lai, G. F. Gao, and W. Shi. The emergence, genomic diversity and global spread of SARS-CoV-2. Nature, 600(7889):408–418, Dec 2021. doi: 10.1038/s41586-021-04188-6. URL https://doi.org/10.1038/s41586-021-04188-6.

N. Ly-Trong, S. Naser-Khdour, R. Lanfear, and B. Q. Minh. AliSim: A Fast and Versatile Phylogenetic Sequence Simulator for the Genomic Era. Molecular Biology and Evolution, 39(5):msac092, 05 2022. doi: 10.1093/molbev/msac092. URL https://doi.org/10.1093/molbev/msac092.

N. Ly-Trong, G. M. J. Barca, and B. Q. Minh. AliSim-HPC: parallel sequence simulator for phylogenetics. Bioinformatics, 39(9):btad540, 09 2023. doi: 10.1093/bioinformatics/btad540. URL https://doi.org/10.1093/bioinformatics/btad540.

K. A. Lythgoe, M. Hall, L. Ferretti, M. de Cesare, G. MacIntyre-Cockett, A. Trebes, M. Andersson, N. Otecko, E. L. Wise, N. Moore, et al. SARS-CoV-2 within-host diversity and transmission. Science, 372(6539):eabg0821, 2021.

B. Q. Minh, H. A. Schmidt, O. Chernomor, D. Schrempf, M. D. Woodhams, A. von Haeseler, and R. Lanfear. IQ-TREE 2: New Models and Efficient Methods for Phylogenetic Inference in the Genomic Era. Molecular Biology and Evolution, 37(5):1530–1534, 02 2020. doi: 10.1093/molbev/msaa015. URL https://doi.org/10.1093/molbev/msaa015.

B. Morel, P. Barbera, L. Czech, B. Bettisworth, L. Hübner, S. Lutteropp, D. Serdari, E.-G. Kostaki, I. Mamais, A. M. Kozlov, P. Pavlidis, D. Paraskevis, and A. Stamatakis. Phylogenetic Analysis of SARS-CoV-2 Data Is Difficult. Molecular Biology and Evolution, 38(5):1777–1791, 12 2020. doi: 10.1093/molbev/msaa314. URL https://doi.org/10.1093/molbev/msaa314.

S. Naser-Khdour, B. Q. Minh, and R. Lanfear. The influence of model violation on phylogenetic inference: a simulation study. bioRxiv, pages 2021–09, 2021.

L. Nesterenko, L. Blassel, P. Veber, B. Boussau, and L. Jacob. Phyloformer: Fast, accurate, and versatile phylogenetic reconstruction with deep neural networks. Molecular Biology and Evolution, 42(4):msaf051, 2025.

L.-T. Nguyen, H. A. Schmidt, A. von Haeseler, and B. Q. Minh. IQ-TREE: A Fast and Effective Stochastic Algorithm for Estimating Maximum-Likelihood Phylogenies. Molecular Biology and Evolution, 32(1):268–274, 11 2014. doi: 10.1093/molbev/msu300. URL https://doi.org/10.1093/molbev/msu300.

B. Pickering, O. Lung, F. Maguire, P. Kruczkiewicz, J. D. Kotwa, T. Buchanan, M. Gagnier,J. L. Guthrie, C. M. Jardine, A. Marchand-Austin, et al. Divergent SARS-CoV-2 variant emerges in white-tailed deer with deer-to-human transmission. Nature Microbiology, 7 (12):2011–2024, 2022.

M. N. Price, P. S. Dehal, and A. P. Arkin. FastTree 2 – Approximately Maximum-Likelihood Trees for Large Alignments. PLOS ONE, 5(3):1–10, 03 2010. doi: 10.1371/journal.pone.0009490. URL https://doi.org/10.1371/journal.pone.0009490.

B. Rannala and Z. Yang. Probability distribution of molecular evolutionary trees: A new method of phylogenetic inference. Journal of Molecular Evolution, 43(3):304–311, Sep 1996. doi: 10.1007/BF02338839. URL https://doi.org/10.1007/BF02338839.

B. Rannala and Z. Yang. Phylogenetic Inference Using Whole Genomes. Annual Review of Genomics and Human Genetics, (Volume 9, 2008):217–231, 2008. doi: 10.1146/annurev.genom.9.081307.164407. URL https://www.annualreviews.org/content/journals/10.1146/annurev.genom.9.081307.164407.

V. Ranwez and O. Gascuel. Quartet-Based Phylogenetic Inference: Improvements and Limits. Molecular Biology and Evolution, 18(6):1103–1116, 06 2001. doi: 10.1093/oxfordjournals.molbev.a003881. URL https://doi.org/10.1093/oxfordjournals.molbev.a003881.

R. M. Rao, J. Liu, R. Verkuil, J. Meier, J. Canny, P. Abbeel, T. Sercu, and A. Rives. MSA Transformer. In M. Meila and T. Zhang, editors, Proceedings of the 38th International Conference on Machine Learning, volume 139 of Proceedings of Machine Learning Research, pages 8844–8856. PMLR, 18–24 Jul 2021. URL https://proceedings.mlr.press/v139/rao21a.html.

D. F. Robinson and L. R. Foulds. Comparison of phylogenetic trees. Mathematical Biosciences, 53(1–2): 131–147, 1981.

F. Ronquist, M. Teslenko, P. van der Mark, D. L. Ayres, A. Darling, S. Höhna, B. Larget,L. Liu, M. A. Suchard, and J. P. Huelsenbeck. MrBayes 3.2: Efficient Bayesian Phylogenetic Inference and Model Choice Across a Large Model Space. Systematic Biology, 61 (3):539–542, 02 2012. doi: 10.1093/sysbio/sys029. URL https://doi.org/10.1093/sysbio/sys029.

N. Saitou and M. Nei. The neighbor-joining method: a new method for reconstructing phylogenetic trees. Molecular Biology and Evolution, 4(4):406–425, 07 1987. doi: 10.1093/oxfordjournals.molbev.a040454. URL https://doi.org/10.1093/oxfordjournals.molbev.a040454.

M. Schuster and K. K. Paliwal. Bidirectional recurrent neural networks. IEEE Transactions on Signal Processing, 45(11): 2673–2681, 1997.

T. Shao, W. Wang, M. Duan, J. Pan, Z. Xin, B. Liu, F. Zhou, and G. Wang. Application of Bayesian phylogenetic inference modelling for evolutionary genetic analysis and dynamic changes in 2019-nCoV. Briefings in Bioinformatics, 22(2): 896–904, 2021.

M. L. Smith and M. W. Hahn. Phylogenetic inference using generative adversarial networks. Bioinformatics, 39(9):btad543, 2023.

R. R. Sokal and C. D. Michener. A statistical method for evaluating systematic relationships. University of Kansas science bulletin, 38: 1409–1438, 1958. URL https://api.semanticscholar.org/CorpusID:61950873.

T. Stadler. Simulating Trees with a Fixed Number of Extant Species. Systematic Biology, 60 (5):676–684, 04 2011. doi: 10.1093/sysbio/syr029. URL https://doi.org/10.1093/sysbio/syr029.

J. G. Sumner, P. D. Jarvis, J. Fernández-Sánchez, B. T. Kaine, M. D. Woodhams, and B. R. Holland. Is the General Time-Reversible Model Bad for Molecular Phylogenetics? Systematic Biology, 61(6):1069–1074, 03 2012. doi: 10.1093/sysbio/sys042. URL https://doi.org/10.1093/sysbio/sys042.

D. L. Swofford and J. Sullivan. Phylogeny inference based on parsimony and other methods using PAUP, page 267–312. Cambridge University Press, 2009.

E. Talevich, B. M. Invergo, P. J. Cock, and B. A. Chapman. Bio.Phylo: A unified toolkit for processing, analyzing and visualizing phylogenetic trees in Biopython. BMC Bioinformatics, 13(1):209, Aug 2012. doi: 10.1186/1471-2105-13-209. URL https://doi.org/10.1186/1471-2105-13-209.

S. Tavaré. Some probabilistic and statistical problems on the analysis of DNA sequence. Lecture of Mathematics for Life Science, 17:57, 1986.

A. Vaswani, N. Shazeer, N. Parmar, J. Uszkoreit, L. Jones, A. N. Gomez, L. u. Kaiser, and I. Polosukhin. Attention is All you Need. 30, 2017. URL https://proceedings.neurips.cc/paper_files/paper/2017/file/3f5ee243547dee91fbd053c1c4a845aa-Paper.pdf.

M. Waterman and T. Smith. On the similarity of dendrograms. Journal of Theoretical Biology, 73(4): 789–800, 1978. doi: 10.1016/0022-5193(78)90137-6. URL https://www.sciencedirect.com/science/article/pii/0022519378901376.

R. J. Williams. Simple statistical gradient-following algorithms for connectionist reinforcement learning. Machine Learning, 8(3):229–256, May 1992. doi: 10.1007/BF00992696. URL https://doi.org/10.1007/BF00992696.

T. Xie and C. Zhang. ARTree: A Deep Autoregressive Model for Phylogenetic Inference. 36: 14427–14444, 2023. URL https://proceedings.neurips.cc/paper_files/paper/2023/file/2e9e513860b1342f3a12ebecf0528a21-Paper-Conference.pdf.

Z. Yang. Maximum likelihood phylogenetic estimation from DNA sequences with variable rates over sites: Approximate methods. Journal of Molecular Evolution, 39(3):306–314, Sep 1994. doi: 10.1007/BF00160154. URL https://doi.org/10.1007/BF00160154.

C. Zhang. Learnable Topological Features For Phylogenetic Inference via Graph Neural Networks. In The Eleventh International Conference on Learning Representations, 2023. URL https://openreview.net/forum?id=hVVUY7p64WL.

M. Zhou, Z. Yan, E. Layne, N. Malkin, D. Zhang, M. Jain, M. Blanchette, and Y. Bengio. PhyloGFN: Phylogenetic inference with generative flow networks. arXiv preprint arXiv:2310.08774, 2023.

X. Zhou, X.-X. Shen, C. T. Hittinger, and A. Rokas. Evaluating Fast Maximum Likelihood-Based Phylogenetic Programs Using Empirical Phylogenomic Data Sets. Molecular Biology and Evolution, 35(2):486–503, 11 2017. doi: 10.1093/molbev/msx302. URL https://doi.org/10.1093/molbev/msx302.

Z. Zou, H. Zhang, Y. Guan, and J. Zhang. Deep Residual Neural Networks Resolve Quartet Molecular Phylogenies. Molecular Biology and Evolution, 37(5):1495–1507, 12 2019. doi: 10.1093/molbev/msz307. URL https://doi.org/10.1093/molbev/msz307.

