## Supplementary for "Accurate and efficient phylogenetic inference through end-to-end deep learning"

[*]Corresponding authors

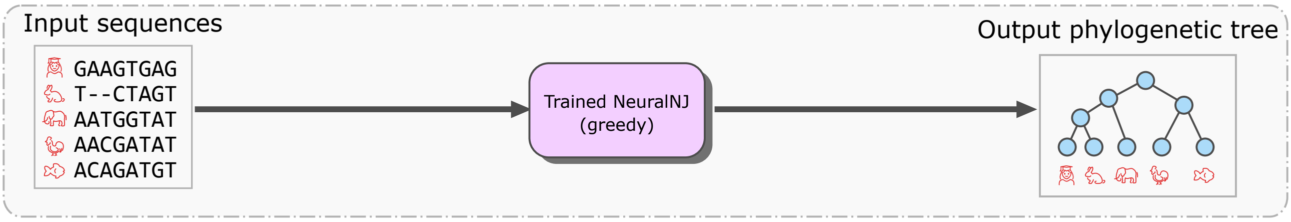


**(a)** NeuralNJ

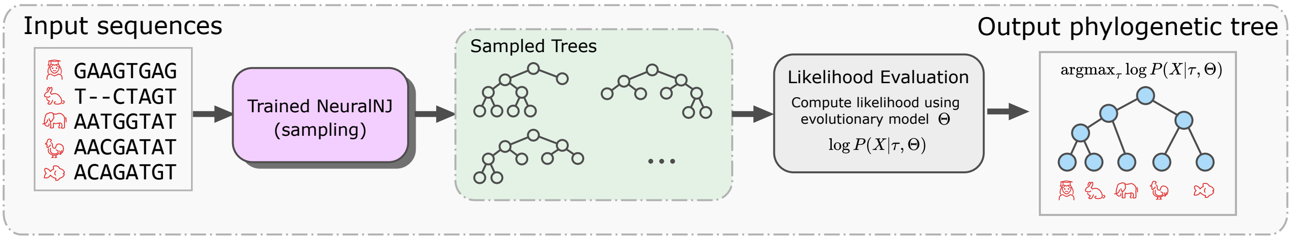


**(b)** NeuralNJ-MC

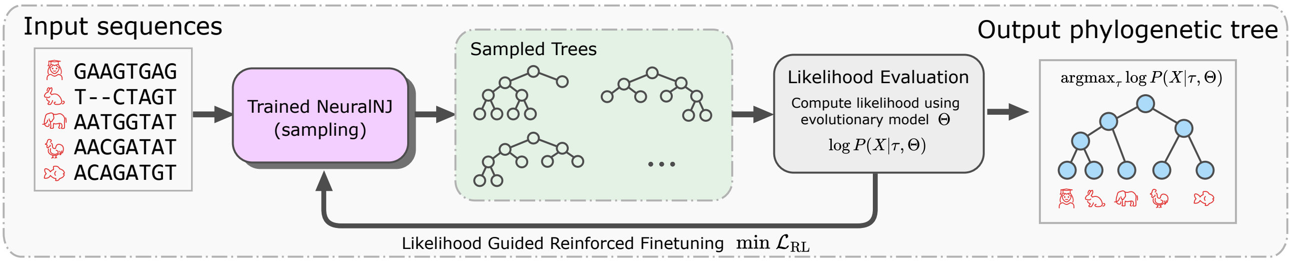


**(c)** NeuralNJ-RL

**Figure S1.** Three inference strategies of NeuralNJ: NeuralNJ, NeuralNJ-MC, and NeuralNJ-RL.

#### A. Three variants of NeuralNJ

Figure S1 illustrates the three inference strategies of NeuralNJ. NeuralNJ employs greedy decoding for direct tree construction, while NeuralNJ-MC extends this by incorporating Monte Carlo sampling (Hastings, 1970) to explore multiple candidate topologies and select the one with highest likelihood. NeuralNJ-RL further enhances the framework through likelihood-guided reinforcement learning for adaptive model refinement. These variants share the same neural architecture but differ in their inference mechanisms, offering a trade-off between computational efficiency and phylogenetic accuracy.

#### B. Implementation details.

All model parameters described below were determined through preliminary experiments to optimize performance.

***B.1. Supervised learning configuration.*** For supervised learning, we trained our models using the Adam optimizer (Kingma and Ba, 2017) with an initial learning rate of 0.0003 and a decay rate of 0.98 per epoch. To monitor training progress, we created a validation set that matched the sequence length distribution of the training set (i.e., 256, 512, and 1,024 nucleotides), ensuring independence from both training and test datasets. Validation performance was checked every 10,000 training steps, and training was terminated when no improvement was observed for five consecutive validation checks (early stopping). Based on this criterion, the optimal model weights were selected at 360,000 training steps (corresponding to epoch 12).

For data generated under the JC evolution model, we configured the model with an embedding dimension of 32, four axial attention layers in the encoder, and four attention heads per layer. For the more complex GTR+I+G evolution model, we increased the configuration to an embedding dimension of 64, six axial attention layers, and eight attention heads per layer. We used a margin of 0.5 for the loss function. The batch size was set to 1 for the simulated dataset. In our network architecture, we handle gaps ('-' characters) by encoding them using a one-hot vector representation that assigns equal probability to all possible bases.

***B.2. Evaluation metrics implementation.*** In addition to the topological accuracy evaluation using RF distance in the main text, we also assessed the phylogenetic inference quality using likelihood-based metrics. The log-likelihood gap is defined as the difference between the log-likelihood scores of RAxML-NG's maximum likelihood tree and those of trees inferred by each method. Unless otherwise specified, all likelihood calculations were performed using the Newton-Raphson method (Felsenstein, 1981), with branch lengths optimized via libpll (Flouri et al., 2014). For completeness, we also analyzed each method's performance using their native branch length estimates, with results presented in Table 5.

***B.3. Selective penalization strategy.*** As mentioned in the main text, we implement a selective penalization strategy that focuses on high-scoring incorrect candidates. This section details the technical implementation of this approach.

Let $\mathcal{C}_t^{\tau_{\text{gt}}} \subset \mathcal{A}_t$ denote the subset of candidate subtrees at step $t$ that are consistent with the ground-truth tree $\tau_{\text{gt}}$. The complement set $\mathcal{A}_t \setminus \mathcal{C}_t^{\tau_{\text{gt}}}$ represents all incorrect candidate subtrees at this step.

For each construction step $t$, we dynamically determine the proportion of incorrect candidates to consider based on training progress:

$$ratio = \max(1 - \alpha \cdot epoch, \beta) \cdot r_f, \tag{S1}$$

where $\alpha = 0.01875$ is the decay rate, $\beta = 0.25$ is the minimum ratio threshold, and $r_f = 0.5$ is a scaling factor which are empirically determined through preliminary experiments on our 50-taxon dataset. This formula ensures that as training progresses, the proportion of considered incorrect candidates decreases from approximately 50% initially to a minimum of 12.5%, allowing the model to gradually focus on more challenging cases while maintaining training stability.

We then select the top $K$ highest scoring incorrect candidates from $\mathcal{A}_t \setminus \mathcal{C}_t^{\tau_{\text{gt}}}$:

$$K = \min(|\mathcal{A}_t \setminus \mathcal{C}_t^{\tau_{\text{gt}}}|, \max(|\mathcal{A}_t \setminus \mathcal{C}_t^{\tau_{\text{gt}}}| \cdot ratio, 8)) \tag{S2}$$

This approach guarantees that at least 8 challenging incorrect candidates are always considered, even in later training stages. Only these $K$ selected highest-scoring incorrect candidates are paired with correct candidates to compute the margin-based pairwise ranking loss described in the main text.

***B.4. Inference and reinforcement learning.*** For phylogenetic inference, NeuralNJ performs direct one-step inference using argmax of priority scores, while NeuralNJ-MC and NeuralNJ-RL employ an iterative search strategy with 100 steps, a parameter empirically determined to balance inference accuracy and computational efficiency (Supplementary Figure S14 and Figure S13). For both NeuralNJ-MC and NeuralNJ-RL, we set the softmax temperature parameter $\gamma$ to 1.0. For reinforcement learning fine-tuning, we employed 10 training epochs with a learning rate of 0.0001. Each training iteration consisted of 10 episodes and 1 baseline episode, with gradient clipping at 0.1 and entropy regularization strength at 0.2. The batch sizes were dynamically adjusted based on the problem scale: 64 for 20 species, 10 for 50 species, and 3 for 100 species at sequence length 1024. For the 100-taxa scenarios with varying sequence lengths, we used batch sizes of 5 and 10 for lengths 256 and 512, respectively.

***B.5. Hardware and software environment.*** All experiments were conducted on an A800 GPU. The implementation was developed using Python 3.9.18 and PyTorch 2.2.1 with CUDA 12.1 support.

**Algorithm 1:** Neural Neighbor Join Algorithm.

**Input:** $n$ aligned sequences data $X = \{x_i\}_{i=1}^{N}$

Embed sequences: $H = \{h_i\}_{i=1}^{N} \leftarrow \mathcal{M}_{\text{Enc}}\left(\{x_i\}_{i=1}^{N}\right)$

Initialize subtrees set with all leave nodes: $S_0 \leftarrow \{\tau_i\}_{i=1}^{N}$

**for** step $t \in [1, \ldots, N-2]$ **do**

  Obtain candidate joined subtrees: $\mathcal{A}_t \leftarrow \{\tau_x = \text{JoinTree}(\tau_i, \tau_j) | \tau_i \in S, \tau_j \in S, i < j\}$

  **for** $\tau_x = \text{JoinTree}(\tau_i, \tau_j)$ `in` $\mathcal{A}_t$ **do**

    **if** priority $s_{\tau_x}$ hasn't been computed **then**

      Aggregate representation for $\tau_x$: $h_x \leftarrow \mathcal{M}_{\text{Est}}\left(h_i, h_j | H, S\right)$

      Compute priority score for $\tau_x$: $s_{\tau_x} \leftarrow \mathcal{M}_{\text{Eval}}\left(h_x\right)$

  Select one joined subtree: $\tau_x^* \sim \pi\left(\tau_x | [s_{\tau_x}]_{\tau_x \in \mathcal{A}_t}\right)$

  $S_t \leftarrow$ Add $\tau_x^*$ into $S_{t-1}$ and remove its children

**return** $\text{JoinTree}(S[0], S[1])$

**Output:** Constructed phylogenetic tree $\tau$

#### C. Algorithm details

This section provides algorithm pseudocode related to NeuralNJ. Algorithm 1 presents the main procedure for phylogenetic tree construction from MSA. The core components include sequence encoder (Algorithm 2), topology-aware parent node representation estimator (Algorithm 3), and subtree evaluation (Algorithm 4). The training procedures for supervised learning and reinforcement learning are detailed in Algorithms 5 and 6, respectively.

**Algorithm 2:** Sequence Encoder.

**function** $\text{SequenceEncoder}(X = [x_i]_{i=1}^{N})$ ▷ $x_i \in \mathbb{R}^L \times \mathbb{R}^4$

  $\tilde{x}_{ij} \leftarrow \text{MLP}\left(x_{ij}\right)$ ▷ $x_i' \in \mathbb{R}^L \times \mathbb{R}^d$

  $\tilde{x}_{ij}' \leftarrow \text{AxialTransformer}\left(\tilde{x}_{ij}\right)$

  $h_{ij} \leftarrow \text{MLP}\left(\tilde{x}_{ij}'\right)$

  **return** $H = \{h_i\}_{i=1}^{N}$

**Algorithm 3:** Parent Node Representation Estimator.

**function** $\text{ParentNodeRepresentationEstimator}(h_l, h_r, H = [h_i])$ ▷ $h_l, h_r, h_i \in \mathbb{R}^{\tilde{L}} \times \mathbb{R}^d$

  $z_j \leftarrow \text{sigmoid}\left(\text{Linear}\left(h_{lj} - h_{rj}\right)\right)$ ▷ Stage 1: Immediate Descendant Integration

  $\tilde{h}_{xj} \leftarrow h_{lj} \odot z_j + h_{jr} \odot (1 - z_j)$

  $q \leftarrow \text{Flatten}\left(\text{Linear}_q\left(h_x\right)\right)$ ▷ Stage 2: Ancestral Context Integration

  $k_i \leftarrow \text{Flatten}\left(\text{Linear}_k\left(h_i\right)\right)$

  $\alpha_i \leftarrow \text{softmax}_i\left(\left[\frac{q \cdot k_i}{\sqrt{\tilde{L} \times d}}, i \neq l \text{ and } i \neq r\right]\right)$ ▷ Context representation through cross-attention

  $h_{\text{rest}} \leftarrow \sum_{i \neq l, r} \alpha_i h_i$

  $w_j \leftarrow \text{sigmoid}\left(\text{Linear}\left(h_{\text{rest}j}\right)\right)$

  $h_{xj} \leftarrow h_{xj} \odot (1 - w_j) + h_{\text{rest}j} \odot w_j$

  **return** $h_x = [h_{xj}]_{j=1}^{\tilde{L}}$

#### D. Additional experimental results

***D.1. Performance evaluation on Jukes-Cantor model data.*** To complement our main analysis under the GTR+I+G model, we also evaluated NeuralNJ's performance on data simulated under the simpler Jukes-Cantor (JC) model (Jukes et al., 1969). The JC model assumes uniform substitution rates and equal base frequencies for DNA sequences, providing a fundamental baseline for phylogenetic inference evaluation (Erixon et al., 2003). Following the experimental framework described in main text section , we trained and evaluated NeuralNJ using identical dataset configurations and baseline methods.

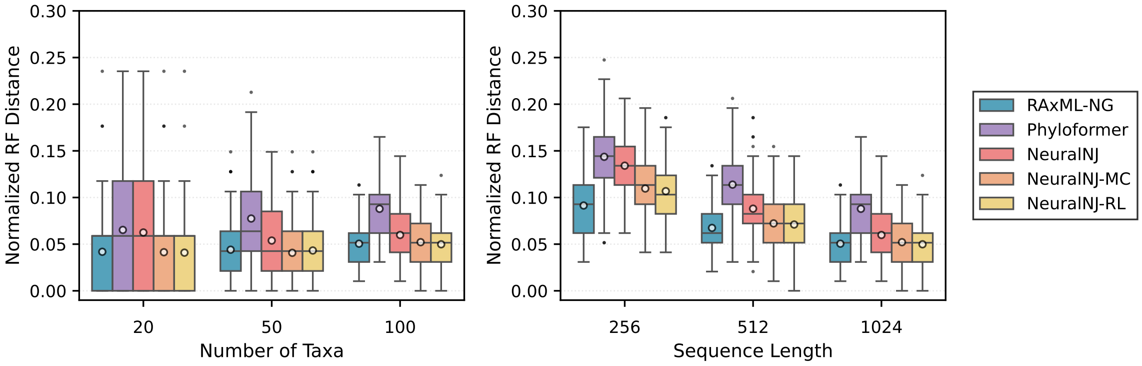


**Figure S2.** Normalized Robinson-Foulds distance comparison of different phylogenetic inference methods across different taxonomic scales and sequence lengths under the JC model. Left: RF distances for varying number of species (n = 20, 50, 100) with fixed sequence length (L = 1024). Right: RF distances for varying sequence lengths (L = 256, 512, 1024) with fixed number of species (n = 100). Box plots show the median (central line), quartiles (box), 1.5× interquartile range (whiskers), outliers (individual points), and the mean value (circle) for each group, with lower RF distances indicating higher accuracy in topology reconstruction.

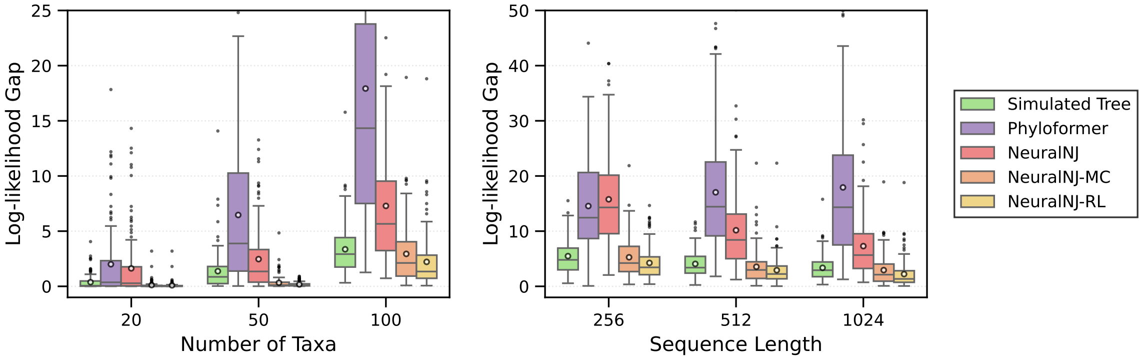


**Figure S3.** Log-likelihood gap comparison between different phylogenetic inference methods and RAxML-NG under the JC model. Left: Results for varying species numbers (n = 20, 50, 100) with fixed sequence length (L = 1024). Right: Results for varying sequence lengths (L = 256, 512, 1024) with fixed species number (n = 100). Box plots show the distribution across 128 replicates, with smaller gaps indicating closer alignment to RAxML-NG's likelihood values. The "Simulated Tree" represents the log-likelihood gap between the true tree (used in simulation) and RAxML-NG's maximum likelihood tree for the corresponding MSA.

**Algorithm 4:** Subtree Evaluator.

**function** $\mathrm{SubtreeEvaluator}(h_x)$
- $s_{\tau_x} \leftarrow \frac{1}{L}\sum_{j=1}^{L}\mathrm{MLP}(h_{xj})$ ▷ Average over sites
- **return** $s_{\tau_x}$

**Algorithm 5:** Supervised Training Algorithm for NeuralNJ.

**function** $\mathrm{Train}$(dataset $\mathcal{D}$, sequence encoder $\mathcal{M}_{\mathsf{Enc}}$, priority predictor $\mathcal{M}_{\mathsf{Eval}}$, representation aggregator $\mathcal{M}_{\mathsf{Est}}$, model parameters $\theta$, number of epochs $E$, learning rate $\alpha$, margin $m$)
- **for** epoch in $[1,...,E]$ **do**
  - **for** $(X,\tau_{\mathrm{gt}}) \sim \mathcal{D}$ **do**
    - $[a_t],[\mathcal{C}_t^{\tau_{\mathrm{gt}}}] \leftarrow \mathrm{SampleTrajectory}\,(\tau_{\mathrm{gt}}, \mathrm{length}\,(X))$
    - $S_0 \leftarrow \{\tau_i\}_{i=1}^{n}$
    - $H = \{h_i\}_{i=1}^{N} \leftarrow \mathcal{M}_{\mathsf{Enc}}\,(X)$
    - $\mathcal{L}_{\mathrm{SL}} \leftarrow 0$
    - **for** step $t \in [1,...,N-2]$ **do**
      - $\mathcal{A}_t \leftarrow \{\tau_x = \mathrm{JoinTree}(\tau_i,\tau_j) | \tau_i \in S, \tau_j \in S, i<j\}$
      - $\mathcal{P}_t \leftarrow \{(\tau_a,\tau_b) | \tau_a \in \mathcal{C}_t^{\tau_{\mathrm{gt}}}, \tau_b \in \mathcal{A}_t \setminus \mathcal{C}_t^{\tau_{\mathrm{gt}}}\}$
      - **for** $\tau_x = \mathrm{JoinTree}(\tau_i,\tau_j)$ `in` $\mathcal{A}_t$ **do**
        - **if** priority $s_{\tau_x}$ hasn't been computed **then**
          - $h_x \leftarrow \mathcal{M}_{\mathsf{Est}}\,(h_i,h_j|H,S)$
          - $s_{\tau_x} \leftarrow \mathcal{M}_{\mathsf{Eval}}\,(h_x)$
      - $\mathcal{L}_{\mathrm{SL}} \leftarrow \mathcal{L}_{\mathrm{SL}} - \frac{1}{|\mathcal{P}_t|}\sum_{(\tau_a,\tau_b)\in\mathcal{P}_t} \max(s_{\tau_a} - s_{\tau_b} - m, 0)$
      - select joined subtree: $\tau_x^* \leftarrow a_t$
      - $S_t \leftarrow$ Add $\tau_x^*$ into $S_{t-1}$ and remove its children
    - $\mathcal{L}_{\mathrm{SL}} \leftarrow \frac{\mathcal{L}_{\mathrm{SL}}}{N-2}$
    - $\theta \leftarrow \theta - \alpha\nabla_\theta\mathcal{L}_{\mathrm{SL}}$

**function** $\mathrm{SampleTrajectory}$(phylogenetic tree $\tau$, taxa number $N$)
- neighbor_set_list $\leftarrow$ [ ]
- action_list $\leftarrow$ [ ]
- **for** step $t \in [1,...,N]$ **do**
  - neighbor_set $\leftarrow \{n \in \tau$.nodes() $|$ $n$.left.is_leaf(), $n$.right.is_leaf()$\}$
  - sampled_subtree $\leftarrow$ random.choice(neighbor_set)
  - action $\leftarrow$ (sampled_subtree.left, sampled_subtree.right)
  - neighbor_set_list $\leftarrow$ neighbor_set_list || [neighbor_set]
  - action_list $\leftarrow$ action_list || [action]
  - $\tau$.remove(sampled_subtree.left)
  - $\tau$.remove(sampled_subtree.right)
- **return** action_list, neighbor_set_list

Figures S2 and S3 present the performance evaluation results under the JC model. For topological accuracy (Figure S2), NeuralNJ variants demonstrate strong performance across different taxonomic scales and sequence lengths. NeuralNJ significantly outperforms Phyloformer in most configurations, while NeuralNJ-MC and NeuralNJ-RL achieve accuracy levels comparable to RAxML-NG.

The log-likelihood analysis (Figure S3) shows that NeuralNJ variants achieve competitive likelihood scores. NeuralNJ maintains smaller likelihood gaps compared to Phyloformer, while NeuralNJ-MC and NeuralNJ-RL achieve likelihood scores approaching those of the simulated ground truth trees and RAxML-NG's maximum likelihood solutions.

**Algorithm 6:** Reinforced Search Algorithm for NeuralNJ.

**function** ReinforcedSearch(MSA of instance $X$, encoder $\mathcal{M}_{\text{Enc}}$, decoder $\mathcal{M}_{\text{Eval}}$, aggregator $\mathcal{M}_{\text{Est}}$, model parameters $\theta$, number of epochs $E$, learning rate $\alpha$, temperature $\gamma$, batch size $B$)
    **for** epoch in $[1, ..., E]$ **do**
        $\mathcal{L}_{\text{RL}} \leftarrow 0$
        **for** batch in $[1, ..., B]$ **do**
            $S_0 \leftarrow \{\tau_i\}_{i=1}^{n}$
            $H = \{h_i\}_{i=1}^{N} \leftarrow \mathcal{M}_{\text{Enc}}\left(\{x_i\}_{i=1}^{N}\right)$ $\mathcal{L}_{\text{RL}} \leftarrow 0$
            log_probs $\leftarrow$ [ ]
            **for** step $t \in [1, ..., N-2]$ **do**
                $\mathcal{A}_t \leftarrow \{\tau_x = \text{JoinTree}(\tau_i, \tau_j) | \tau_i \in S, \tau_j \in S, i < j\}$
                **for** $\tau_x = \text{JoinTree}(\tau_i, \tau_j)$ `in` $\mathcal{A}_t$ **do**
                    **if** priority $s_{\tau_x}$ hasn't been computed **then**
                        $h_x \leftarrow \mathcal{M}_{\text{Est}}(h_i, h_j | H, S)$
                        $s_{\tau_x} \leftarrow \mathcal{M}_{\text{Eval}}(h_x)$
                logits $\leftarrow \text{LogSoftmax}\left(\left[\frac{s_{\tau_x}}{\gamma}\right]\right)$
                $\tau_x^* \sim \text{Multinomial}\left(\tau_x | [s_{\tau_x}]_{\tau_x \in \mathcal{A}_t}\right)$
                log_probs $\leftarrow$ log_probs $\|$ [logits$[\tau_x^*]$]
                $S_t \leftarrow$ Add $\tau_x^*$ into $S_{t-1}$ and remove its children
            $\hat{\tau} \leftarrow \text{JoinTree}(S[0], S[1])$
            $G \leftarrow \text{TopologyEvaluator}(\hat{\tau}, X)$ ▷ Calculate optimize goal, i.e., likelihood $P(X|\tau)$
            $$\mathcal{L}_{\text{RL}} \leftarrow \mathcal{L}_{\text{RL}} + \left(-\left(\sum_{t=1}^{N-2} \text{log\_probs}[t]\right) \cdot (G - b)\right)$$
        $$\mathcal{L}_{\text{RL}} \leftarrow \frac{1}{B}\mathcal{L}_{\text{RL}}$$
        $\theta \leftarrow \theta - \alpha \nabla_\theta \mathcal{L}_{\text{RL}}$

These results under the JC model demonstrate that NeuralNJ maintains strong performance even with simplified evolutionary assumptions. The consistent performance across both JC and GTR+I+G models (presented in the main text) suggests the robustness of our approach across different evolutionary scenarios.

***D.2. Quantity analysis of performance on simulated data under the JC model.*** To supplement Figure S2 and Figure S3, we present detailed quantitative comparisons under the JC model across varying experimental conditions. Table 1 evaluate performance across taxa numbers (n = 20, 50, 100), while Table 2 evaluates performance across sequence lengths (L = 256, 512, 1024). The detailed metrics show that NeuralNJ outperforms existing learning-based methods, while NeuralNJ-MC and NeuralNJ-RL achieve performance comparable to RAxML-NG in both likelihood scores and topological accuracy (RF distance), particularly for larger phylogenetic problems.

***D.3. Quantity analysis of performance on simulated data under the GTR+I+G model.*** To supplement the topological accuracy results presented in the main text (Figure 2), we evaluated the likelihood performance of different methods (Figure S4). As shown in Figure S4, all NeuralNJ variants maintain likelihood scores comparable to ground truth trees, confirming their effectiveness.

Table 3 and Table 4 provide detailed quantitative comparisons under the GTR+I+G model across taxa numbers (n = 20-100) and sequence lengths (L = 256-1024), respectively. The metrics in Table 3 and Table 4 demonstrate that NeuralNJ outperforms both Phyloformer and RAxML-NG in topology reconstruction, while NeuralNJ-MC and NeuralNJ-RL further enhance accuracy through likelihood-guided search.

To examine the impact of branch length optimization, Table 5 to compare the effect of branch length optimization using libpll on likelihood estimation for Phyloformer and Simulated Tree methods. The results suggest that the absence of branch length optimization increases the log-likelihood gap, but the relative ranking of methods in terms of topological accuracy remains unchanged. This demonstrates that our main conclusions regarding topology reconstruction are robust to the choice of branch length optimization strategy.

***D.4. Analysis on computational efficiency and accuracy under the JC model.*** We conducted similar efficiency-accuracy trade-off analyses under the JC model (Figures S5 and S6). The results show consistent patterns with those ob-

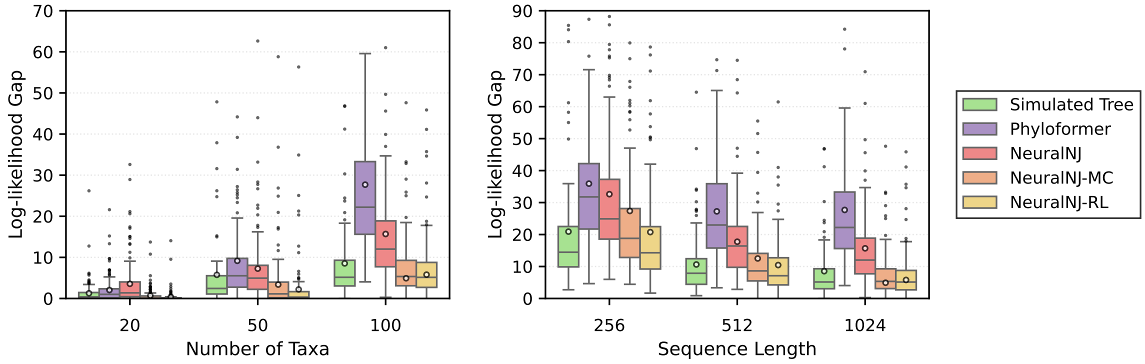


**Figure S4.** Log-likelihood gaps between each predicted tree and the RAxML-NG baseline under GTR+I+G across varying numbers of taxa (left) and sequence lengths (right). Smaller gaps indicate better phylogenetic inference. Box plots show the median (central line), quartiles (box), 1.5× interquartile range (whiskers), outliers (individual points), and the mean value (circle) for each group.

**Table 1.** NeuralNJ (ours) v.s. baselines on synthetic task sets under the JC model across different task sizes. LLH Gap represents the log-likelihood difference from RAxML-NG, and RF denotes the Robinson-Foulds distance with the simulated tree. Lower values indicate better performance (↓). Best results are shown in **bold**, with superscripts $^1$, $^2$, and $^3$ indicating the first, second, and third best performances respectively.

| Method | $n = 20$ | | $n = 50$ | | $n = 100$ | |
|---|---|---|---|---|---|---|
| | LLH Gap ↓ | RF ↓ | LLH Gap ↓ | RF ↓ | LLH Gap ↓ | RF ↓ |
| Simulated Tree | 0.379 | **0.000**$^1$ | 1.371 | **0.000**$^1$ | 3.357 | **0.000**$^1$ |
| BIONJ (Gascuel, 1997) | 44.236 | 0.179 | 284.363 | 0.284 | 858.016 | 0.348 |
| PhyML (Guindon et al., 2010) | **0.012**$^2$ | 0.044 | 2.803 | 0.045 | 3.254 | 0.052 |
| RAxML-NG (Kozlov et al., 2019) | **0.000**$^1$ | 0.042 | **0.000**$^1$ | 0.044 | **0.000**$^1$ | **0.050**$^2$ |
| Phyloformer (Nesterenko et al., 2025) | 1.999 | 0.065 | 6.465 | 0.077 | 17.926 | 0.088 |
| **NeuralNJ (ours)** | 1.626 | 0.062 | 2.465 | 0.054 | 7.289 | 0.060 |
| **NeuralNJ-MC (ours)** | 0.111 | 0.041$^3$ | 0.323$^3$ | **0.041**$^2$ | 2.940$^3$ | 0.052 |
| **NeuralNJ-RL (ours)** | 0.091$^3$ | **0.041**$^2$ | **0.176**$^2$ | 0.043$^3$ | **2.218**$^2$ | **0.050**$^2$ |

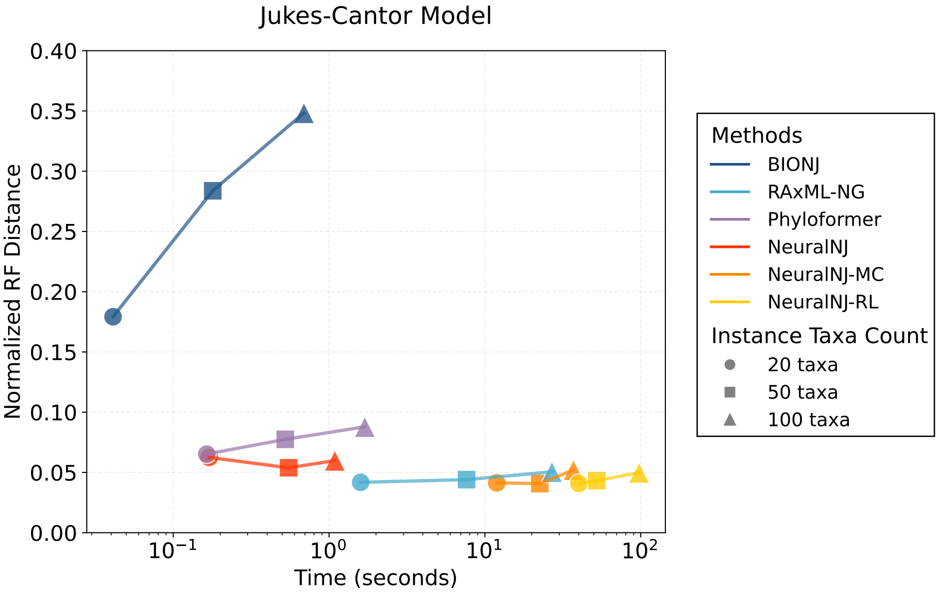


**Figure S5.** Performance comparison of phylogenetic inference methods on simulated data under JC models with varying taxonomic scales (n = 20, 50, 100) and fixed sequence length (L = 1024). The x-axis represents computational time in seconds (log scale), and the y-axis shows the normalized Robinson-Foulds distance. Different colors indicate different methods, while markers denote varying numbers of taxa (circles: 20, squares: 50, triangles: 100).

served under the GTR+I+G model (presented in the main text): BIONJ achieves the fastest computation but with limited accuracy, while NeuralNJ variants demonstrate balanced performance across different scales. Specifically, NeuralNJ maintains efficiency comparable to Phyloformer while achieving better accuracy, and NeuralNJ-MC/RL variants achieve accuracy levels close to RAxML-NG with moderate computational overhead. The consistent performance patterns across both evolutionary models further validate the scalability and efficiency of our approach.

***D.5. Detailed efficiency-accuracy analysis across problem scales under GTR+I+G model.*** Figures S7 and S8 provide detailed performance trajectories for each method as problem scales vary. With increasing taxa numbers (Figure S7), these methods show degraded accuracy, but with distinct patterns: BIONJ's accuracy deteriorates rapidly despite maintaining fast computation, while RAxML-NG achieves better accuracy but with substantial computational overhead. NeuralNJ variants demonstrate more graceful scaling, with NeuralNJ maintaining efficient computation while achieving accuracy comparable to slower methods.

For varying sequence lengths (Figure S8), the performance trajectories reveal method-specific scaling characteristics. NeuralNJ and Phyloformer maintain consistent computation times across sequence lengths due to their parallel processing architecture. In contrast, RAxML-NG exhibit increased computational costs with sequence length,

**Table 2.** NeuralNJ (ours) v.s. baselines on simulated task sets under JC model across different sequence lengths. LLH Gap represents the log-likelihood difference from RAxML-NG, and RF denotes the Robinson-Foulds distance with the simulated tree. Lower values indicate better performance (↓). Best results are shown in **bold**, with superscripts $^1$, $^2$, and $^3$ indicating the first, second, and third best performances respectively.

| Method | $L = 256$ | | $L = 512$ | | $L = 1024$ | |
|---|---|---|---|---|---|---|
| | LLH Gap ↓ | RF ↓ | LLH Gap ↓ | RF ↓ | LLH Gap ↓ | RF ↓ |
| Simulated Tree | 5.478 | **0.000**$^1$ | 4.073 | **0.000**$^1$ | 3.357 | **0.000**$^1$ |
| BIONJ (Gascuel, 1997) | 248.938 | 0.337 | 529.395 | 0.382 | 858.016 | 0.348 |
| PhyML (Guindon et al., 2010) | **1.572**$^2$ | 0.093$^3$ | 3.476$^3$ | 0.071$^3$ | 3.254 | 0.052 |
| RAxML-NG (Kozlov et al., 2019) | **0.000**$^1$ | **0.091**$^2$ | **0.000**$^1$ | **0.067**$^2$ | **0.000**$^1$ | **0.050**$^2$ |
| Phyloformer (Nesterenko et al., 2025) | 14.555 | 0.144 | 17.038 | 0.114 | 17.926 | 0.088 |
| **NeuralNJ (ours)** | 15.772 | 0.134 | 10.165 | 0.088 | 7.289 | 0.060 |
| **NeuralNJ-MC (ours)** | 5.263 | 0.110 | 3.546 | 0.072 | 2.940$^3$ | 0.052 |
| **NeuralNJ-RL (ours)** | 4.242$^3$ | 0.107 | **2.921**$^2$ | 0.071$^3$ | **2.218**$^2$ | **0.050**$^2$ |

**Table 3.** NeuralNJ (ours) v.s. baselines on synthetic task sets under GTR+I+G evolution model across different task sizes. LLH Gap represents the log-likelihood difference from RAxML-NG, and RF denotes the Robinson-Foulds distance with the simulated tree. Lower values indicate better performance (↓). Best results are shown in **bold**, with superscripts $^1$, $^2$, and $^3$ indicating the first, second, and third best performances respectively.

| Method | $n = 20$ | | $n = 50$ | | $n = 100$ | |
|---|---|---|---|---|---|---|
| | LLH Gap ↓ | RF ↓ | LLH Gap ↓ | RF ↓ | LLH Gap ↓ | RF ↓ |
| Simulated Tree | 1.268 | **0.000**$^1$ | 5.752 | **0.000**$^1$ | 8.535 | **0.000**$^1$ |
| RAxML-NG (Kozlov et al., 2019) | **0.000**$^1$ | 0.108 | **0.000**$^1$ | 0.155 | **0.000**$^1$ | 0.142 |
| Phyloformer (Nesterenko et al., 2025) | 2.058 | 0.116 | 9.162 | 0.133 | 27.689 | 0.159 |
| **NeuralNJ (ours)** | 3.566 | 0.114 | 7.262 | 0.104 | 15.686 | 0.112$^3$ |
| **NeuralNJ-MC (ours)** | 0.699$^3$ | 0.089$^3$ | 3.384$^3$ | **0.096**$^2$ | **4.922**$^2$ | **0.102**$^2$ |
| **NeuralNJ-RL (ours)** | **0.384**$^2$ | **0.087**$^2$ | **2.199**$^2$ | 0.102$^3$ | 5.797$^3$ | **0.102**$^2$ |

**Table 4.** NeuralNJ (ours) v.s. baselines on simulated task sets of GTR+I+G evolution model across different sequence lengths. LLH Gap represents the log-likelihood difference from RAxML-NG, and RF denotes the Robinson-Foulds distance with the simulated tree. Lower values indicate better performance (↓). Best results are shown in **bold**, with superscripts $^1$, $^2$, and $^3$ indicating the first, second, and third best performances respectively.

| Method | $L = 256$ | | $L = 512$ | | $L = 1024$ | |
|---|---|---|---|---|---|---|
| | LLH Gap ↓ | RF ↓ | LLH Gap ↓ | RF ↓ | LLH Gap ↓ | RF ↓ |
| Simulated Tree | 20.913$^3$ | **0.000**$^1$ | 10.633$^3$ | **0.000**$^1$ | 8.535 | **0.000**$^1$ |
| RAxML-NG (Kozlov et al., 2019) | **0.000**$^1$ | 0.316 | **0.000**$^1$ | 0.206 | **0.000**$^1$ | 0.142 |
| Phyloformer (Nesterenko et al., 2025) | 35.918 | 0.300 | 27.244 | 0.215 | 27.689 | 0.159 |
| **NeuralNJ (ours)** | 32.624 | 0.269 | 17.752 | 0.171 | 15.686 | 0.112$^3$ |
| **NeuralNJ-MC (ours)** | 27.372 | 0.268$^3$ | 12.517 | 0.163$^3$ | **4.922**$^2$ | **0.102**$^2$ |
| **NeuralNJ-RL (ours)** | **20.761**$^2$ | **0.262**$^2$ | **10.450**$^2$ | **0.161**$^2$ | 5.797$^3$ | **0.102**$^2$ |

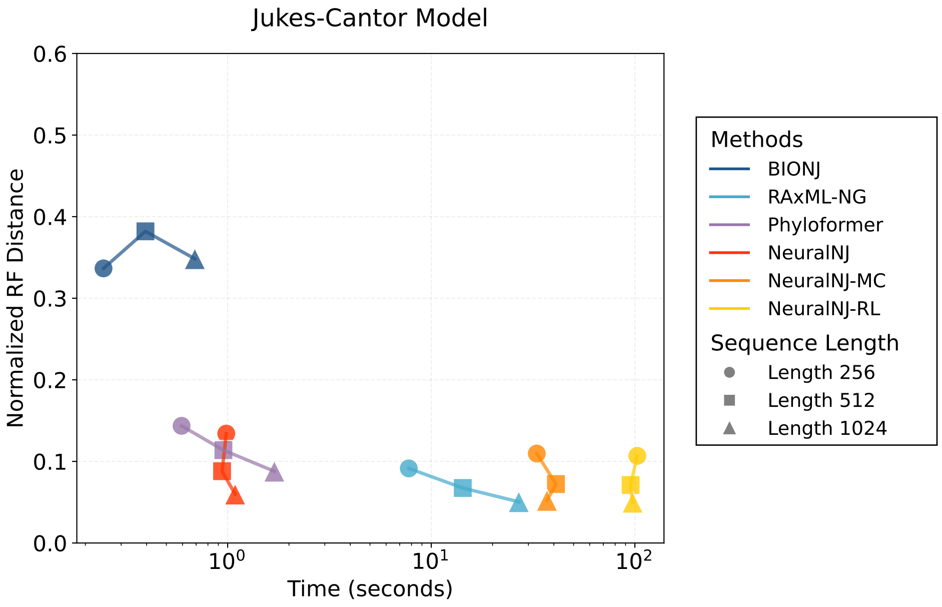


**Figure S6.** Performance comparison of phylogenetic inference methods on simulated data under JC models with fixed number of taxa (n = 100) and varying sequence lengths. The x-axis represents computational time in seconds (log scale), and the y-axis shows the normalized Robinson-Foulds distance. Different colors indicate different methods, while markers denote sequence lengths (circles: 256, squares: 512, triangles: 1024 nucleotides).

due to its iterative likelihood optimizations. Notably, all methods show improved accuracy with increasing sequence

**Table 5.** Comparison of Mean Log-likelihood Gap with RAxML-NG for different methods on test datasets with varying taxa counts ($n$) and sequence lengths ($L$) without branch length optimization. Methods labeled with "(w/o brlen opt)" indicate results without branch length optimization using libpll. Lower values indicate better performance.

| Method | $n = 20$ | $n = 50$ | $n = 100$ | $L = 256$ | $L = 512$ | $L = 1024$ |
|---|---|---|---|---|---|---|
| Simulated Tree | 1.268 | 5.752 | 8.535 | 20.913 | 10.633 | 8.535 |
| Simulated Tree(w/o brlen opt) | 1.235 | 7.944 | 21.688 | 23.821 | 11.421 | 21.688 |
| Phyloformer | 2.058 | 9.162 | 27.689 | 35.918 | 27.244 | 27.689 |
| Phyloformer(w/o brlen opt) | 2.142 | 12.958 | 37.436 | 38.749 | 27.822 | 37.436 |
| **NeuralNJ (ours)** | 3.566 | 7.262 | 15.686 | 32.624 | 17.752 | 15.686 |
| **NeuralNJ-MC (ours)** | 0.699 | 3.384 | 4.922 | 27.372 | 12.517 | 4.922 |
| **NeuralNJ-RL (ours)** | 0.384 | 2.199 | 5.797 | 20.761 | 10.450 | 5.797 |

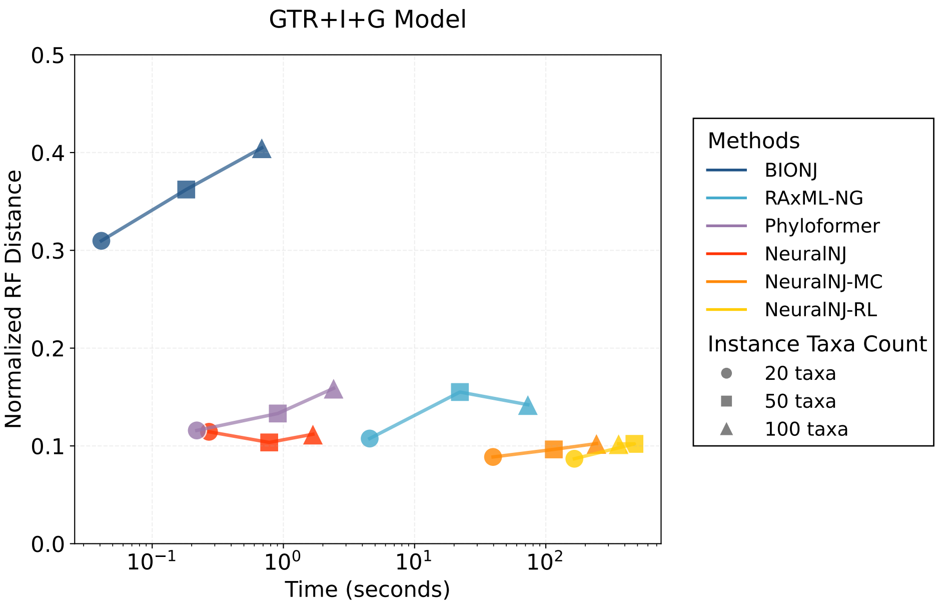


**Figure S7.** Performance comparison of phylogenetic inference methods on simulated data under GTR+I+G models with varying taxonomic scales (n = 20, 50, 100) and fixed sequence length (L = 1024). The x-axis represents computational time in seconds (log scale), and the y-axis shows the normalized Robinson-Foulds distance. Different colors indicate different methods, while markers denote varying numbers of taxa (circles: 20, squares: 50, triangles: 100).

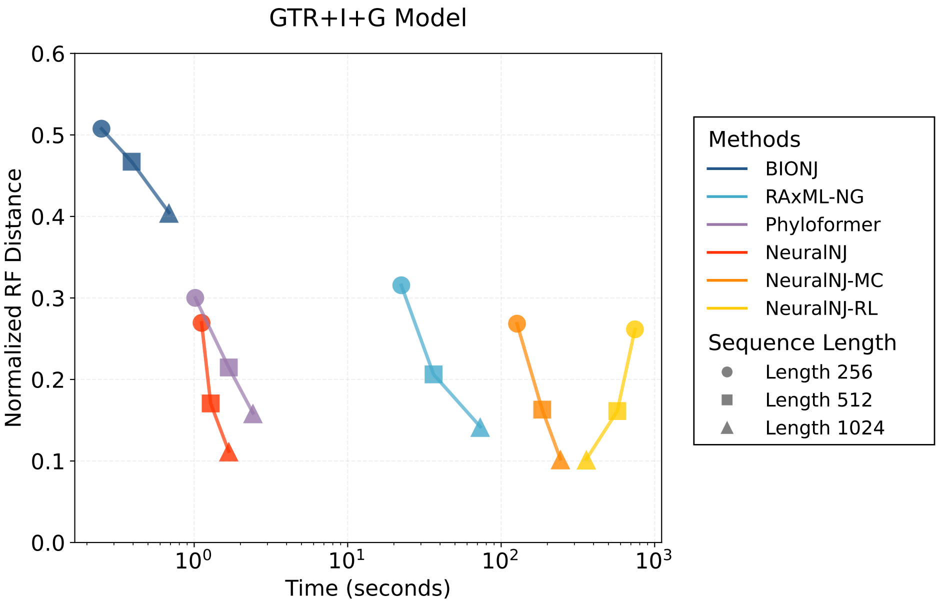


**Figure S8.** Performance comparison of phylogenetic inference methods on simulated data under GTR+I+G models with fixed number of taxa (n = 100) and varying sequence lengths. The x-axis represents computational time in seconds (log scale), and the y-axis shows the normalized Robinson-Foulds distance. Different colors indicate different methods, while markers denote sequence lengths (circles: 256, squares: 512, triangles: 1024 nucleotides).

length, with NeuralNJ variants achieving the best accuracy-efficiency trade-off. These scaling behaviors explain the Pareto-optimal performance of NeuralNJ variants observed in the main analysis.

The consistent performance of NeuralNJ across both dimensions (taxa count and sequence length) validates its design choices: priority-based tree construction and parallel sequence processing enable efficient scaling, while the neural architecture maintains accuracy across problem scales. These detailed trajectories complement the main text's Pareto analysis by revealing the underlying scaling mechanisms that contribute to NeuralNJ's balanced performance.

***D.6. Time on simulated data under the JC and GTR+I+G model.*** To complement the efficiency analysis (Figure S5, Figure S6 Figure S7 and Figure S8), Table 6 and Table 7 present detailed runtime comparisons across different experimental settings for both JC and GTR+I+G models. NeuralNJ demonstrates comparable efficiency to Phyloformer, completing inference within 1-2 seconds for most scenarios. While NeuralNJ-MC and NeuralNJ-RL require additional computation time due to their iterative search strategies, these variants offer a trade-off between computational cost and phylogenetic inference precision. Specifically, for a dataset with 100 taxa and 1024 sites, NeuralNJ completes inference in less 2 seconds under both JC and GTR+I+G models, offering an efficient alternative to traditional methods like RAxML-NG (26.91 and 73.01 seconds, respectively). This analysis underscores the balance between computational efficiency and accuracy offered by the NeuralNJ variants, particularly in large-scale phylogenetic analyses.

**Table 6.** Time (seconds) of NeuralNJ (ours) v.s. baselines on different simulated task sets of JC model (n: number of taxa, L: sequence length).

| Method | $n = 20$ | $n = 50$ | $n = 100$ | $L = 256$ | $L = 512$ | $L = 1024$ |
|---|---|---|---|---|---|---|
| BIONJ (Gascuel, 1997) | 0.04 | 0.18 | 0.69 | 0.25 | 0.39 | 0.69 |
| PhyML (Guindon et al., 2010) | 5.97 | 31.90 | 121.19 | 35.30 | 68.28 | 121.19 |
| RAxML-NG (Kozlov et al., 2019) | 1.59 | 7.63 | 26.91 | 7.76 | 14.28 | 26.91 |
| Phyloformer (Nesterenko et al., 2025) | 0.16 | 0.52 | 1.70 | 0.59 | 0.95 | 1.70 |
| **NeuralNJ (ours)** | 0.17 | 0.55 | 1.09 | 0.98 | 0.94 | 1.09 |
| **NeuralNJ-MC (ours)** | 11.92 | 22.51 | 36.99 | 32.99 | 40.97 | 36.99 |
| **NeuralNJ-RL (ours)** | 40.05 | 52.22 | 97.51 | 102.78 | 95.31 | 97.51 |

**Table 7.** Time (seconds) of NeuralNJ (ours) v.s. baselines on different simulated task sets of GTR+I+G evolution model (n: number of taxa, L: sequence length).

| Method | $n = 20$ | $n = 50$ | $n = 100$ | $L = 256$ | $L = 512$ | $L = 1024$ |
|---|---|---|---|---|---|---|
| BIONJ (Saitou and Nei, 1987) | 0.04 | 0.18 | 0.68 | 0.25 | 0.39 | 0.68 |
| RAxML-NG (Kozlov et al., 2019) | 4.55 | 22.15 | 73.01 | 22.35 | 36.28 | 73.01 |
| Phyloformer (Nesterenko et al., 2025) | 0.22 | 0.91 | 2.41 | 1.02 | 1.68 | 2.41 |
| **NeuralNJ (ours)** | 0.27 | 0.78 | 1.68 | 1.12 | 1.28 | 1.68 |
| **NeuralNJ-MC (ours)** | 39.62 | 115.09 | 243.33 | 126.81 | 185.14 | 243.33 |
| **NeuralNJ-RL (ours)** | 165.13 | 473.70 | 358.93 | 744.26 | 571.05 | 358.93 |

***D.7. Absolute log likelihood values performance for all approaches.*** We report the absolute log-likelihood values for all methods in Tables 8 and 9. These values complement the relative performance metrics (LLH Gap) presented in the main text (Tables 1, 2 for JC model and Tables 3, 4 for GTR+I+G model). The absolute values offer additional context for assessing the statistical fit of inferred phylogenies across different experimental conditions.

**Table 8.** Log-likelihood of NeuralNJ (ours) v.s. baselines on simulated task sets under JC model across different task sizes.

| Method | $n = 20$ | $n = 50$ | $n = 100$ | $L = 256$ | $L = 512$ | $L = 1024$ |
|---|---|---|---|---|---|---|
| BIONJ (Gascuel, 1997) | -12819.166 | -27324.786 | -50572.647 | -16664.123 | -26871.367 | -50572.647 |
| PhyML (Guindon et al., 2010) | -12774.942 | -27043.226 | -49717.885 | -16416.757 | -26345.448 | -49717.885 |
| RAxML-NG (Kozlov et al., 2019) | -12774.930 | -27040.423 | -49714.631 | -16415.185 | -26341.972 | -49714.631 |
| Phyloformer (Nesterenko et al., 2025) | -12776.928 | -27046.888 | -49732.557 | -16429.740 | -26359.010 | -49732.557 |
| **NeuralNJ (ours)** | -12776.555 | -27042.888 | -49721.920 | -16430.957 | -26352.137 | -49721.920 |
| **NeuralNJ-MC (ours)** | -12775.040 | -27040.764 | -49717.571 | -16420.448 | -26345.518 | -49717.571 |
| **NeuralNJ-RL (ours)** | -12775.021 | -27040.599 | -49716.849 | -16419.427 | -26344.893 | -49716.849 |

**Table 9.** Log-likelihood of NeuralNJ (ours) v.s. baselines on different simulated task sets under GTR+I+G evolution model.

| Method | $n = 20$ | $n = 50$ | $n = 100$ | $L = 256$ | $L = 512$ | $L = 1024$ |
|---|---|---|---|---|---|---|
| Simulated Tree | -13088.518 | -28207.669 | -51806.366 | -14208.587 | -27691.737 | -51806.366 |
| RAxML-NG (Kozlov et al., 2019) | -13087.250 | -28201.918 | -51797.832 | -14187.674 | -27681.104 | -51797.832 |
| Phyloformer (Nesterenko et al., 2025) | -13089.309 | -28211.080 | -51825.521 | -14223.592 | -27708.348 | -51825.521 |
| **NeuralNJ (ours)** | -13090.816 | -28209.179 | -51813.517 | -14220.297 | -27698.856 | -51813.517 |
| **NeuralNJ-MC (ours)** | -13087.949 | -28205.302 | -51802.753 | -14215.046 | -27693.621 | -51802.753 |
| **NeuralNJ-RL (ours)** | -13087.634 | -28204.116 | -51803.629 | -14208.435 | -27691.554 | -51803.629 |

***D.8. Additional experimental results on empirical datasets.*** Table 10 reports the mean normalized RF distances for each method on the complete datasets. These results correspond to the quantitative values underlying the box plots in Figure 3 of the main text. The result indicates that NeuralNJ generally outperforms Phyloformer, while RAxML-NG achieves the highest accuracy overall. Notably, NeuralNJ-MC further improves upon NeuralNJ's results in most cases.

We further evaluated performance on the longest feasible sequence alignments within hardware constraints. For JarvD5a, SongD1, and TarvD7, we selected alignments of 10,000–17,000 nucleotides; for WickD3b, we used 3,000–4,000 nucleotides. This yielded subsets of SongD1 (6 cases), WickD3b (5 cases), JarvD5a (348 cases), and TarvD7 (90 cases). Table 11 summarizes the normalized RF distances on these long sequence alignments. As shown, all methods achieve substantially improved accuracy on longer sequences.

In summary, across all empirical evaluations, the relative performance pattern remains consistent: NeuralNJ and its variants generally outperform Phyloformer, while RAxML-NG achieves the highest accuracy overall. NeuralNJ-MC and NeuralNJ-RL further enhance accuracy, especially on sampled datasets, demonstrating the effectiveness and robustness of our sampling-based approaches.

***D.9. Additional results on long simulated data.*** To investigate the performance of NeuralNJ on longer sequences, we conducted experiments with alignment lengths of 2048 and 4096 for 100 taxa. Following the same simulation method used for the shorter sequences (256, 512, and 1024) in the main text, we generated 128 unique test tree-MSA pairs for each sequence length. Due to GPU memory constraints, results for NeuralNJ-RL at lengths 2048 and 4096 are not available, as this variant requires the most memory resources. Figure S9 presents box plots of the normalized Robinson-Foulds (RF) distances for all methods across sequence lengths ranging from 256 to 4096. Figure S10 illustrates the relationship between average inference time per sample and sequence length for each method.

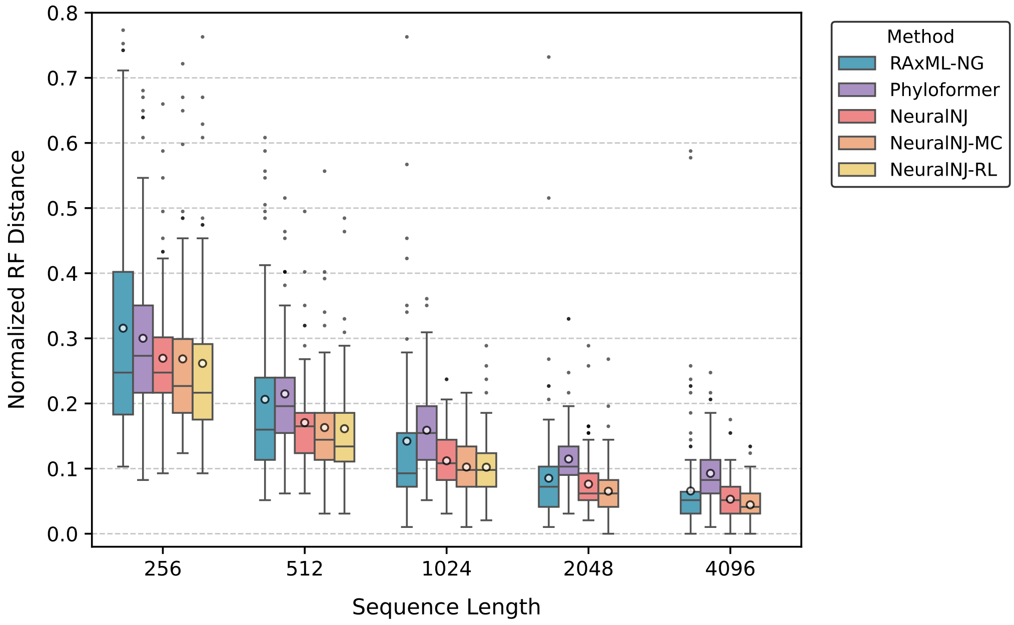


**Figure S9.** Normalized Robinson-Foulds (RF) distances of RAxML-NG, Phyloformer, NeuralNJ, NeuralNJ-MC, and NeuralNJ-RL on data with 100 taxa across sequence lengths from 256 to 4096 base pairs.

As shown in Figure S9, all methods generally exhibit improved accuracy (lower RF distances) as sequence length increases. Beyond the superior performance of NeuralNJ and its variants at shorter lengths (256, 512, and 1024)

**Table 10.** Average normalized RF distances on complete empirical datasets

| Dataset | RAxML | Phyloformer | NeuralNJ | NeuralNJ-MC |
|---|---|---|---|---|
| JarvD5a | 0.743 | 0.780 | 0.778 | 0.690 |
| SongD1 | 0.267 | 0.527 | 0.451 | 0.369 |
| TarvD7 | 0.366 | 0.557 | 0.546 | 0.443 |
| WickD3b | 0.483 | 0.572 | 0.580 | 0.469 |

**Table 11.** Normalized RF distances on long sequence alignments from empirical datasets

| Dataset | RAxML | Phyloformer | NeuralNJ | NeuralNJ-MC |
|---|---|---|---|---|
| JarvD5a | 0.558 | 0.660 | 0.682 | 0.622 |
| SongD1 | 0.152 | 0.412 | 0.348 | 0.309 |
| TarvD7 | 0.213 | 0.504 | 0.542 | 0.535 |
| WickD3b | 0.320 | 0.550 | 0.576 | 0.435 |

discussed in the main text, NeuralNJ maintains excellent accuracy with longer sequences. At a sequence length of 2048, NeuralNJ achieves lower RF distances compared to RAxML-NG and Phyloformer, with NeuralNJ-MC showing further improvement. At the longest sequence length of 4096, NeuralNJ and RAxML-NG perform comparably, while NeuralNJ-MC attains the lowest RF distances among all methods.

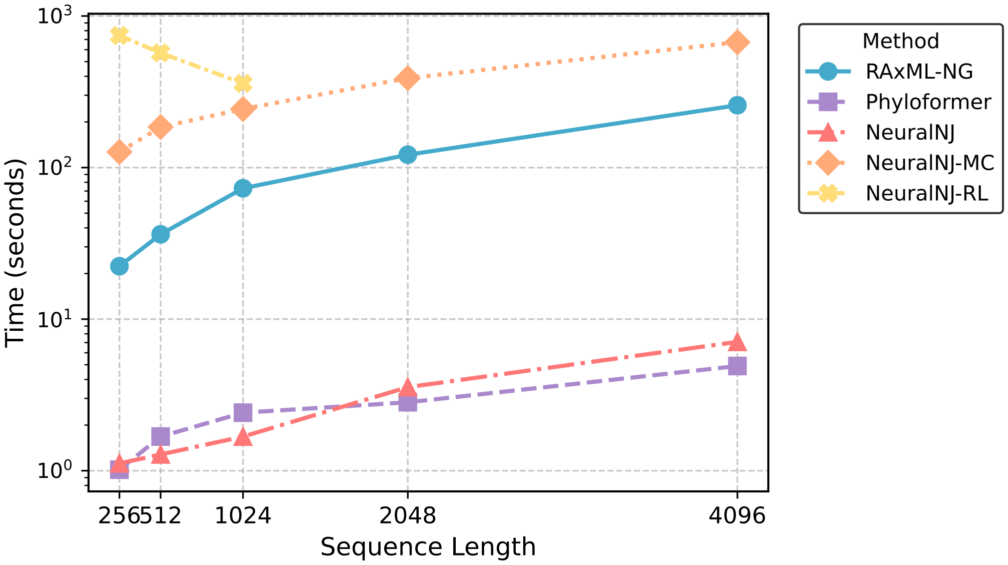


**Figure S10.** Inference time (seconds) of RAxML-NG, Phyloformer, NeuralNJ, NeuralNJ-MC, and NeuralNJ-RL for phylogenetic reconstruction with 100 taxa across sequence lengths from 256 to 4096 base pairs. NeuralNJ and Phyloformer demonstrate 1-2 orders of magnitude faster inference compared to RAxML-NG.

Figure S10 illustrates the computational efficiency of each method. Notably, NeuralNJ and Phyloformer demonstrate inference times 1-2 orders of magnitude faster than the likelihood-based RAxML-NG method. NeuralNJ-MC requires more time due to its additional likelihood calculations, placing it in the same order of magnitude as RAxML-NG. For phylogenetic inference with 100 taxa and 4096 base pairs, both NeuralNJ and Phyloformer complete inference in under 10 seconds, while RAxML-NG requires several minutes. This highlights NeuralNJ's exceptional efficiency.

These results demonstrate that NeuralNJ consistently delivers both high accuracy and high efficiency for long sequence phylogenetic inference.

***D.10. Training and validation curves.*** Figure S11 illustrates the training and validation curves of NeuralNJ. The y-axis represents the mean normalized RF distance, while the x-axis indicates the number of training samples processed (with 29,000 samples constituting one epoch). We present experimental results spanning 410,000 training steps (approximately 14 epochs). The mean RF distance for the training set was calculated across examples with sequence lengths of 256, 512, and 1,024 base pairs. The validation curve reflects performance on an independent validation set featuring 50 taxa and sequence lengths of 256, 512, and 1,024, distinct from both the training and test datasets.

Analysis of the validation curve reveals that the Normalized RF distance reaches its minimum at 360,000 training steps (corresponding to epoch 12), after which the validation performance plateaus with minor fluctuations. Consequently, we selected the model weights from 360,000 training steps as our final model.

***D.11. Analysis of Kuhner-Felsenstein Distances.*** To complement the Robinson-Foulds distance results presented in the main text Figure 2, Figure S12 illustrates the Kuhner-Felsenstein (KF) distances between inferred trees and ground-truth simulated trees for RAxML-NG, Phyloformer, NeuralNJ, NeuralNJ-MC, and NeuralNJ-RL. These comparisons span varying numbers of taxa (20, 50, 100) with fixed sequence length (1024), and varying sequence lengths (256, 512, 1024) with fixed taxon count (100).

KF distance quantifies differences between phylogenetic trees by considering both topological structure and branch lengths, providing providing better quality of trees with branch length than topology-only metrics such as Robinson-Foulds distance. For branch length optimization in the NeuralNJ variants, we employed the libpll software package after establishing tree topologies.

Figure S12 demonstrates that NeuralNJ variants consistently achieve lower KF distances across all experimental conditions. As taxon count increases from 20 to 100, KF distances increase for all methods, yet NeuralNJ-based approaches consistently outperform Phyloformer and RAxML. Notably, at shorter sequence lengths (256 and 512),

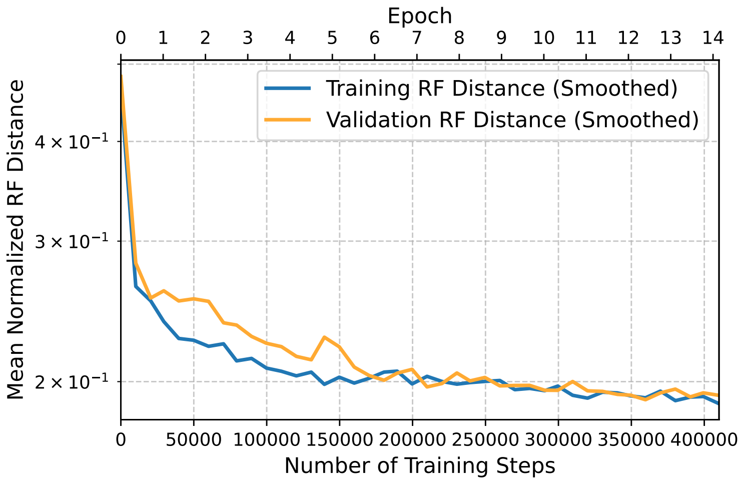


**Figure S11.** Training and validation curves for the NeuralNJ model. The y-axis represents Mean Normalized RF distance while the x-axis shows the number of training samples processed. The top x-axis additionally indicates the corresponding epoch number.

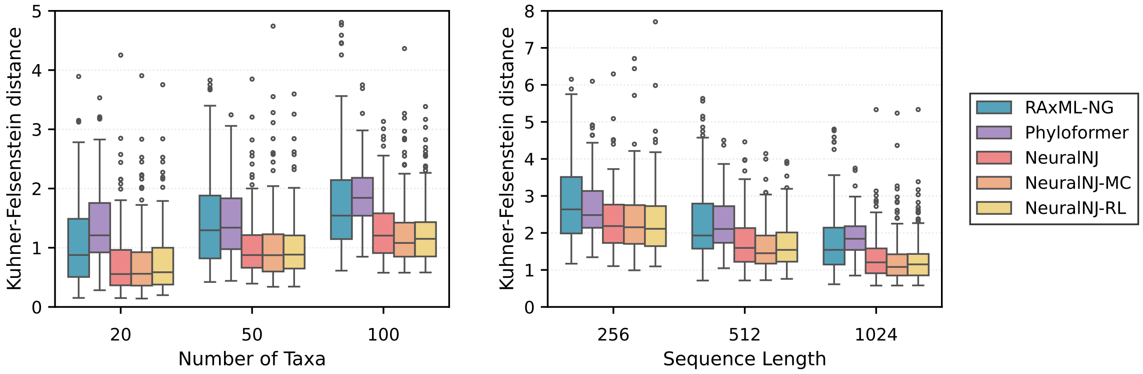


**Figure S12.** Normalized KF distances between predicted and ground-truth simulated trees across varying numbers of taxa (left) and sequence lengths (right). Lower KF distances indicate higher topological accuracy.

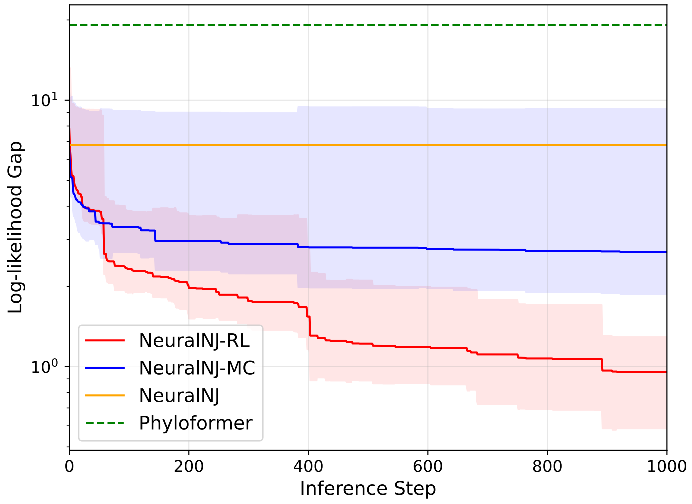


**Figure S13.** Comparison of log-likelihood gaps over search steps between different methods. The plot shows the convergence behavior of three NeuralNJ variants (NeuralNJ-RL, NeuralNJ-MC, and NeuralNJ) against Phyloformer baseline. The shaded areas represent confidence intervals, and lower values indicate better performance.

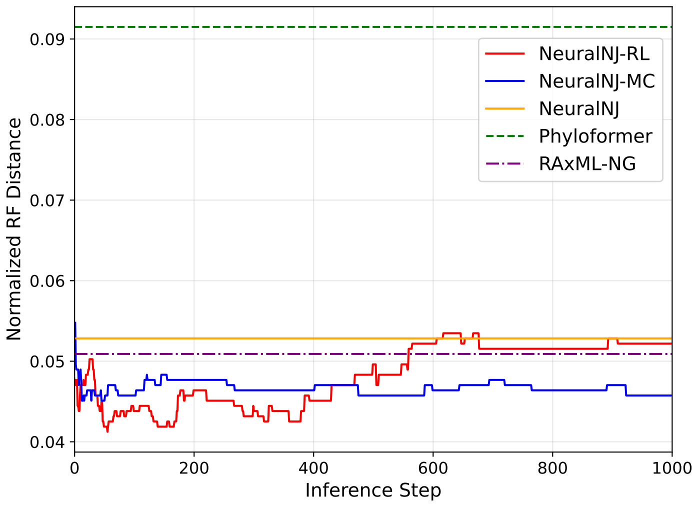


**Figure S14.** Comparison of RF distance over search steps between different methods. The plot shows the convergence behavior of three NeuralNJ variants (NeuralNJ-RL, NeuralNJ-MC, and NeuralNJ) against Phyloformer and RAxML-NG baselines.

NeuralNJ variants exhibit stable performance with minimal variance. As sequence length increases to 1024, NeuralNJ variants further demonstrate performance superiority.

***D.12. Analysis of search strategies.*** To validate the efficacy of our search strategies, we conducted experiments on a subset of simulated data comprising 100 taxa with sequence lengths of 1024 under the JC evolutionary model. We randomly selected 16 test instances for in-depth analysis, each executing 2,000 search steps. The average performance is illustrated in Figure S13 and Figure S14.

Figure S13 demonstrates that all three variants of NeuralNJ outperform the neural network solver Phyloformer. While NeuralNJ employs a greedy approach for rapid solutions, NeuralNJ-MC systematically explores higher likelihood solutions over time. NeuralNJ-RL leverages likelihood optimization for fine-tuning, enabling it to theoretically approach the performance of the professional maximum likelihood solver RAxML-NG.

As shown in Figure S14, both NeuralNJ-MC and NeuralNJ-RL achieve superior topological accuracy compared to Phyloformer, with RF distances consistently lower throughout the search process. NeuralNJ-MC maintains stable performance around 0.045 RF distance, while NeuralNJ-RL exhibits an initial rapid descent followed by gradual convergence. These results demonstrate that our likelihood-guided search strategies effectively optimize tree topology, with NeuralNJ-MC showing particularly stable performance and NeuralNJ-RL demonstrating potential for exploring diverse solution spaces through its adaptive search strategy.

The decreasing log-likelihood gap observed in NeuralNJ-MC and NeuralNJ-RL demonstrates their progressive refinement and effectiveness. Notably, NeuralNJ-RL reduces the gap with RAxML-NG to below 1 as the number of steps increases, highlighting its precision. These findings validate the effectiveness of NeuralNJ's search strategies in exploring a broader solution space and achieving better topologies.

To further clarify the mechanisms of MC/RL search strategies, we present three representative cases in Figure S15, Figure S16, and Figure S17. These cases respectively illustrate scenarios where the initial NeuralNJ solution is worse than, equal to, or better than RAxML-NG in terms of RF distance.

In all cases, the log-likelihood gap decreases as search progresses, reflecting effective likelihood optimization. However, the impact on RF distance varies: in Figure S15, MC/RL strategies substantially reduce RF distance, outperforming both NeuralNJ and RAxML-NG; in Figure S16, MC shows improvement while RL exhibits non-monotonic behavior, ultimately converging to the baseline; in Figure S17, further likelihood optimization by MC/RL may increase RF distance, highlighting a potential trade-off between likelihood and topology.

These representative cases highlight the diverse behaviors of MC and RL search strategies and illustrate that likelihood optimization does not always guarantee improved topological accuracy.

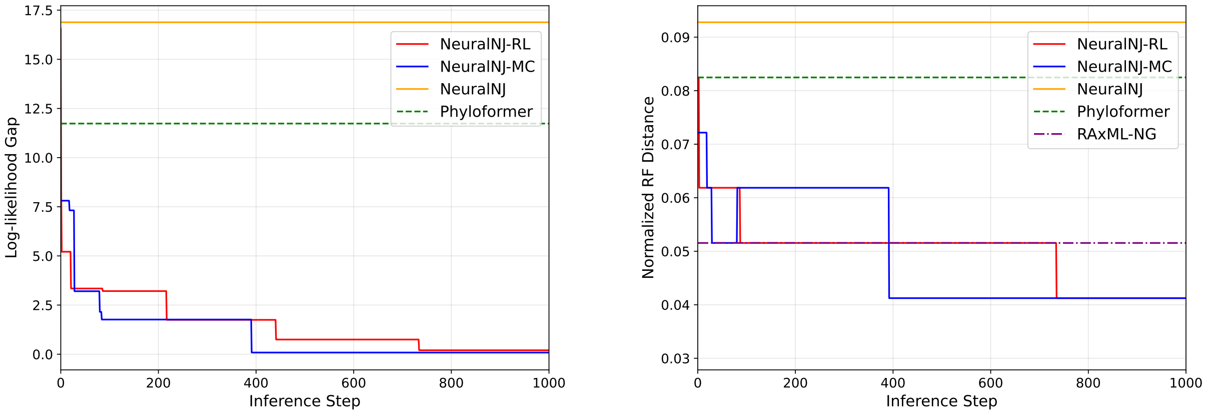


**Figure S15.** Comparison of log-likelihood gap (left) and RF distance (right) over search steps for a representative test instance where the initial NeuralNJ solution has a higher RF distance than RAxML-NG. The plots show the convergence behavior of NeuralNJ-RL and NeuralNJ-MC variants alongside NeuralNJ, Phyloformer, and RAxML-NG baselines.

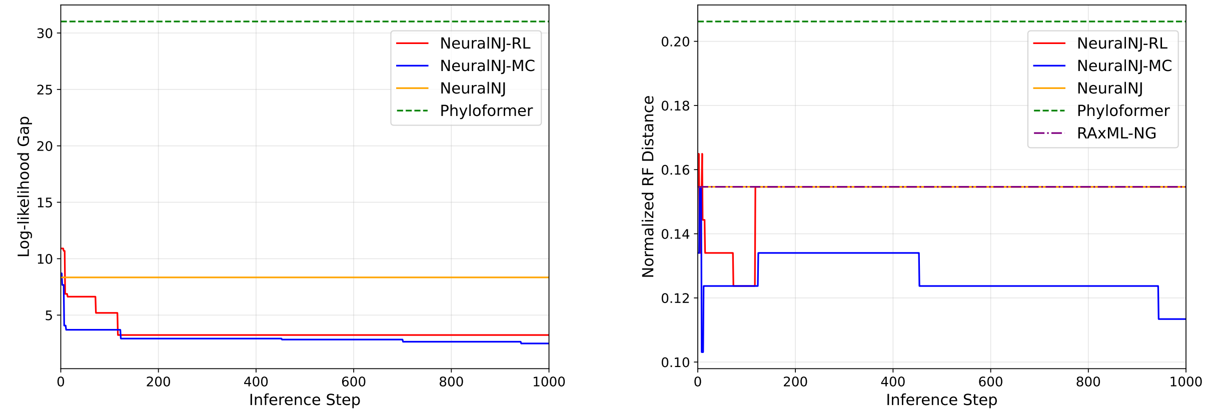


**Figure S16.** Comparison of log-likelihood gap (left) and RF distance (right) over search steps for a representative test instance where the initial NeuralNJ and RAxML-NG solutions have similar RF distances. The plots show the convergence behavior of NeuralNJ-RL and NeuralNJ-MC variants alongside NeuralNJ, Phyloformer, and RAxML-NG baselines.

***D.13. Performance of topology-guided parent node representation estimatorå.*** To validate the effectiveness of our topology-guided parent node representation estimator, we conducted comprehensive ablation experiments. This

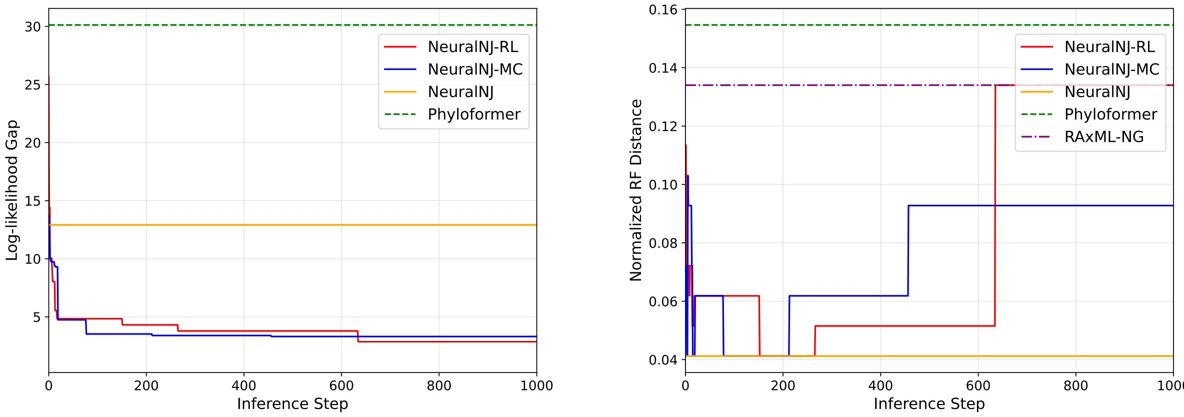


**Figure S17.** Comparison of log-likelihood gap (left) and RF distance (right) over search steps for a representative test instance where the initial NeuralNJ solution has a lower RF distance than RAxML-NG. The plots show the convergence behavior of NeuralNJ-RL and NeuralNJ-MC variants alongside NeuralNJ, Phyloformer, and RAxML-NG baselines.

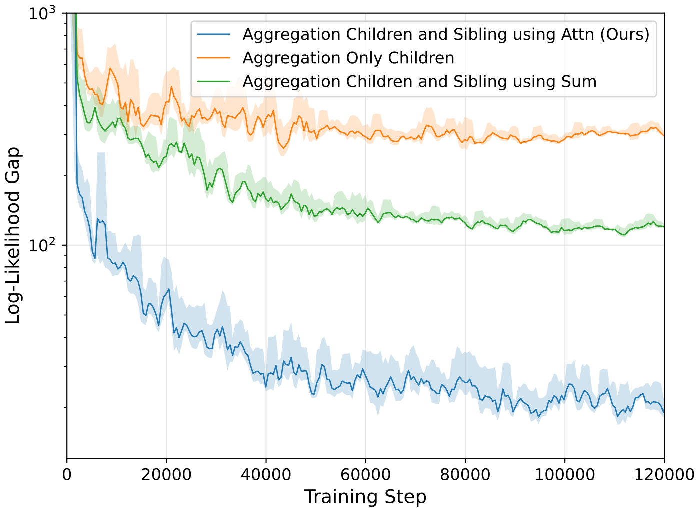


**Figure S18.** Convergence curves on validation set for different representation aggregation techniques.

network models the evolutionary process by which species inherit and diverge from common ancestors, crucial for accurate phylogenetic inference. We compared our full model against two simplified variants to highlight the importance of hierarchical information processing.

Figure S18 compares three aggregation strategies: (1) our complete topology-guided network, which processes child node information and incorporates broader evolutionary context through cross-attention mechanisms; (2) a simplified child-only aggregation; and (3) an intermediate approach using average pooling instead of attention for sibling and ancestral node information. Experiments on the simulated dataset with 50 taxa under the JC model measured performance by likelihood scores during training.

Results show that our topology-guided network achieves superior convergence, maintaining lower likelihood gaps throughout training, with a clear advantage after the initial phase. The child-only strategy shows poorer convergence, while the average-pooling variant performs better but remains suboptimal. These differences align with biological understanding: our model's superior performance reflects its ability to capture both vertical inheritance and broader evolutionary context, essential in molecular evolution. The child-only strategy's limited performance stems from its narrow focus, while the average-pooling variant's intermediate results suggest simple averaging is inadequate for complex relationships. The cross-attention mechanism effectively integrates diverse evolutionary information, underscoring its value in modeling molecular evolution's intricate processes. These findings empirically support our model design and validate its utility for phylogenetic inference.

**Table 12.** Comparison of computational resources between NeuralNJ variants and Phyloformer.

| Method | Inference Mem. (GB) | Training Mem. (GB) | Training Epochs | Training Time (Days) | Parameters (K) |
|---|---|---|---|---|---|
| Phyloformer (Nesterenko et al., 2025) | 3.7 | 63.3 | 19 | 5 | 307.297 |
| **NeuralNJ (ours)** | 3.7 | 12.7 | 12 | 15 | 425.857 |
| **NeuralNJ-MC (ours)** | 3.7 | - | - | - | - |
| **NeuralNJ-RL (ours)** | 11.1 | - | - | - | - |

***D.14. Comparison of computational resources between NeuralNJ variants and Phyloformer.*** In Table 12, we present a comprehensive comparison of computational resource requirements for NeuralNJ variants and Phyloformer during both training and inference phases. All experiments were conducted under the GTR+I+G evolutionary model, with models trained on the simulated dataset comprising various configurations: 50 taxa and sequence lengths of 256, 512, and 1024 nucleotides. For inference evaluation, we utilized the test dataset with 50 taxa and a fixed sequence length of 1024 nucleotides. To ensure fair comparison, both Phyloformer and NeuralNJ models were configured with a batch size of 1 during training and validation phases.

The results reveal several notable patterns. While NeuralNJ demonstrates comparable inference memory usage (3.7 GB) to Phyloformer, it achieves significantly reduced training memory requirements (12.7 GB vs. 63.3 GB). However, this memory efficiency comes at the cost of extended training duration (15 days vs. 5 days) and slightly less training epochs (12 vs. 19). Note that these epoch counts represent the optimal points selected for inference models based on validation performance. The NeuralNJ-MC maintains the same inference memory footprint, while NeuralNJ-RL requires a higher inference memory allocation (11.1 GB), reflecting the additional computational overhead of its learning strategy. Notably, NeuralNJ employs a slightly larger parameter count (425.9K vs. 307.3K), indicating a more complex model architecture while maintaining competitive memory efficiency.

#### E. MSA data used in phylogenetic analysis case study

The ground-truth phylogenetic tree for this case study is illustrated in Figure S19. All genome sequences and corresponding multiple sequence alignments used in our experiments are publicly available in our repository at `https://github.com/DingShizhe/NeuralNJ`.

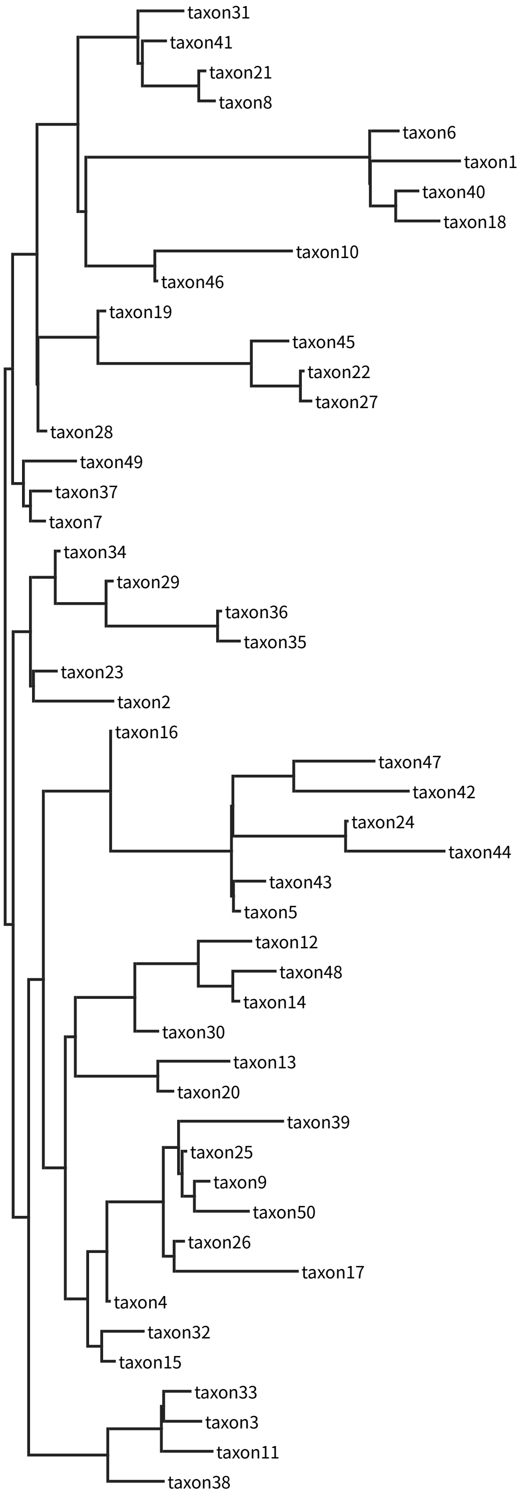


**Figure S19.** The ground truth phylogenetic tree used in the case study. The tree topology was randomly generated, and branch lengths were sampled from a distribution. The MSA data was then simulated based on this tree using sequence evolution models.

#### F. MSA data used in topology construction process analysis

The ground-truth phylogenetic tree for this case study in topology construction process analysis is illustrated in Figure S20. All genome sequences and corresponding multiple sequence alignments used in our experiments are publicly available in our repository at `https://github.com/DingShizhe/NeuralNJ`.

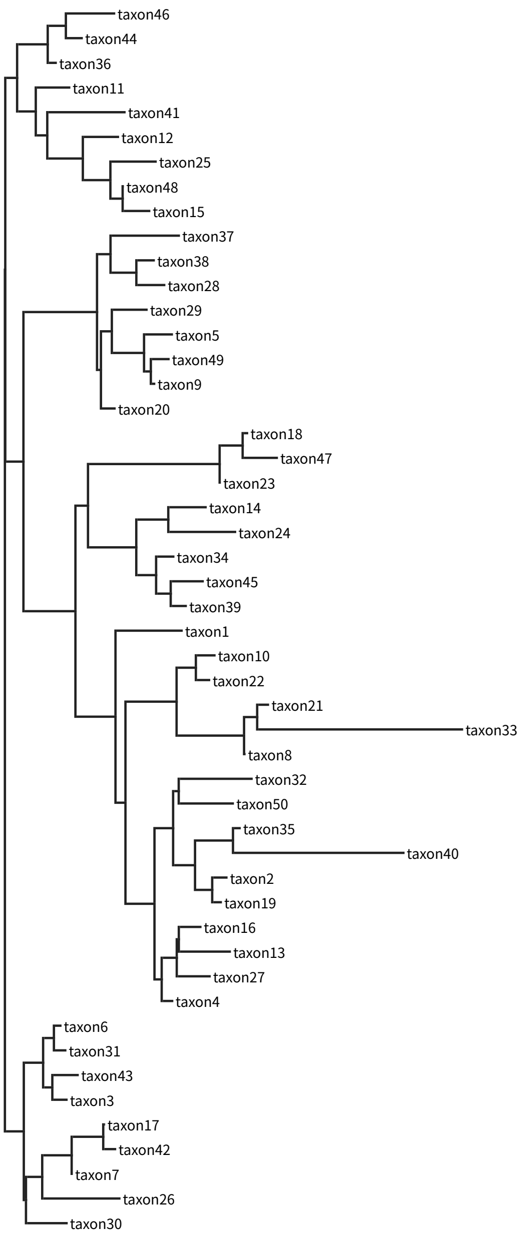


**Figure S20.** The ground truth phylogenetic tree used in the construction process study. The tree topology was randomly generated, and branch lengths were sampled from a distribution. The MSA data was then simulated based on this tree using sequence evolution models.

### Bibliography

P. Erixon, B. Svennblad, T. Britton, and B. Oxelman. Reliability of Bayesian Posterior Probabilities and Bootstrap Frequencies in Phylogenetics. *Systematic Biology*, 52(5):665–673, 10 2003. doi: 10.1080/10635150390235485. URL https://doi.org/10.1080/10635150390235485.

J. Felsenstein. Evolutionary trees from DNA sequences: A maximum likelihood approach. *Journal of Molecular Evolution*, 17(6):368–376, Nov 1981. doi: 10.1007/BF01734359. URL https://doi.org/10.1007/BF01734359.

T. Flouri, F. Izquierdo-Carrasco, D. Darriba, A. Aberer, L.-T. Nguyen, B. Minh, A. Von Haeseler, and A. Stamatakis. The Phylogenetic Likelihood Library. *Systematic Biology*, 64(2):356–362, 10 2014. doi: 10.1093/sysbio/syu084. URL https://doi.org/10.1093/sysbio/syu084.

O. Gascuel. BIONJ: an improved version of the NJ algorithm based on a simple model of sequence data. *Molecular Biology and Evolution*, 14(7):685–695, 1997.

S. Guindon, J.-F. Dufayard, V. Lefort, M. Anisimova, W. Hordijk, and O. Gascuel. New Algorithms and Methods to Estimate Maximum-Likelihood Phylogenies: Assessing the Performance of PhyML 3.0. *Systematic Biology*, 59(3):307–321, 05 2010. doi: 10.1093/sysbio/syq010. URL https://doi.org/10.1093/sysbio/syq010.

W. K. Hastings. Monte Carlo sampling methods using Markov chains and their applications. 1970.

T. H. Jukes, C. R. Cantor, et al. Evolution of protein molecules. *Mammalian Protein Metabolism*, 3(24):21–132, 1969.

D. P. Kingma and J. Ba. Adam: A Method for Stochastic Optimization. 2017. URL https://arxiv.org/abs/1412.6980.

A. M. Kozlov, D. Darriba, T. Flouri, B. Morel, and A. Stamatakis. RAxML-NG: a fast, scalable and user-friendly tool for maximum likelihood phylogenetic inference. *Bioinformatics*, 35(21): 4453–4455, 05 2019. doi: 10.1093/bioinformatics/btz305. URL https://doi.org/10.1093/bioinformatics/btz305.

L. Nesterenko, L. Blassel, P. Veber, B. Boussau, and L. Jacob. Phyloformer: Fast, accurate, and versatile phylogenetic reconstruction with deep neural networks. *Molecular Biology and Evolution*, 42(4):msaf051, 2025.

N. Saitou and M. Nei. The neighbor-joining method: a new method for reconstructing phylogenetic trees. *Molecular Biology and Evolution*, 4(4):406–425, 07 1987. doi: 10.1093/oxfordjournals.molbev.a040454. URL https://doi.org/10.1093/oxfordjournals.molbev.a040454.
